# Genomic profiling of HIV-1 integration in microglia links viral insertions to TAD organization

**DOI:** 10.1101/2022.02.14.480322

**Authors:** Mona Rheinberger, Ana Luisa Costa, Martin Kampmann, Dunja Glavas, Iart Luca Shytaj, Carlotta Penzo, Nadine Tibroni, Oliver T. Fackler, Kristian Vlahovicek, Bojana Lucic, Carl Herrmann, Marina Lusic

**Affiliations:** Department of Infectious Diseases, Integrative Virology, Heidelberg University Hospital, Heidelberg, 69120, Germany; German Center for Infection Research (DZIF), Heidelberg, 69120, Germany; Health Data Science Unit, Medical Faculty University Heidelberg and BioQuant, Heidelberg, 69120, Germany; Bioinformatics Group, Division of Molecular Biology, Department of Biology, Faculty of Science, University of Zagreb, Zagreb, Croatia

## Abstract

HIV-1 persists in anatomically distinct cellular and tissue reservoirs as a stably integrated provirus that is a major barrier to HIV-1 cure. Proviral insertions are largely characterized in blood cells, while HIV-1 integration patterns remain unexplored in microglia, the major brain reservoir. Here, we employ genomics approaches to obtain the first HIV-1 integration site (IS) profiling in microglia and perform in-depth analysis of transcriptome, specific histone signatures and chromatin accessibility on different genomic scales. We show that HIV-1 follows genic insertion patterns into introns of actively transcribed genes, characteristic of blood reservoirs. HIV-1 insertional hotspot analysis by non-negative matrix factorization (NMF)-based approach clusters IS signatures with genic- and super-enhancers. Chromatin accessibility transcription factor (TF) footprints reveal that increased CTCF binding marks latently infected microglia compared to productively infected one. We identify CTCF-enriched topologically associated domain (TAD) borders with signatures of active chromatin as a neighborhood for HIV-1 integration in microglia and CD4^+^ T cells. Our findings further strengthen the notion that HIV-1 follows the patterns of host cell genome organization to integrate and to establish the silent proviral state and reveal that these principles are largely conserved in different anatomical latent reservoirs.

## INTRODUCTION

Human immunodeficiency virus 1 (HIV-1) causes the Acquired Immune Deficiency Syndrome (AIDS) and patients infected with the virus require lifelong antiretroviral therapy (ART), as viral load rebounds from the latent proviral reservoirs when treatment is interrupted (1). The well studied main target cells of HIV-1 are CD4^+^ T cells and macrophages, although latent HIV-1 reservoirs are found throughout the body, including gut-associated lymphoid tissues (GALT), bone marrow and the the central nervous system (CNS), all of which contribute to viral persistence (2–4). Patient data from CNS-derived samples support the existence of replication competent proviruses likely in microglia, years after ART (5–11), It seems likely that low levels of HIV-1 RNA detected in CSF and plasma from suppressed patients contribute to chronic inflammation, and, together with ART toxicity, microglial activation and neuroinflammation lead to HIV-associated neurological disorders (HAND) in people living with HIV (PLWH) (12–18). These clinical implications emphasize the importance of further studies of the main HIV-1 reservoir in the CNS. To this end, patterns of HIV-1 integration and genome regulation linked to viral insertions in microglia still remain undetermined. HIV-1 integration is a non-random process, and IS patterns into transcriptionally active genes have been characterized comprehensively in CD4^+^ T cells and macrophages, major reservoirs of latent provirus (19–30). In this regard, recent studies demonstrated that HIV-1 integration favours speckle-associated domains (SPADs), genomic regions mapped to the gene-dense genome, thus refining HIV-1 targeting biases towards highly active genes (29, 31).

The role of 3D compartmentalization of the T cell genome and vicinity to cell type-specific enhancers termed super-enhancers (SE) was recently ascribed to HIV-1 recurrent integrations (21). Recurrent integration genes (RIGs) and hotspots of integration were found proximal to SE that cluster together in the transcriptionally active A1 subcompartment, pointing towards a structure-function relationship between the 3D genome organization, integration site preferences and the expression of the HIV-1 provirus in T cells (21, 32–35). A and B subcompartments segregate chromosomes based on the long-range interactions enclosed within large (>1MB) gene-active and gene-inactive regions, respectively (33). On a more local scale (<1MB), DNA is folded into chromatin loops and topologically associated domains (TADs) (21, 32–39). TADs are characterized by high frequency of internal DNA interactions that facilitate enhancer-promoter contacts, while their margins, referred to as TAD boundaries, are largely conserved among different cell types and are enriched for the insulator binding protein CCCTC-binding factor (CTCF) (36–39). In the context of HIV-1 infection, a recent study applying Assay for Transposase-Accessible Chromatin using sequencing (ATAC-Seq) to actively and latently infected T cells suggested an increased occupancy and transcriptional repression mediated by architectural protein CTCF in latent cells, thus proposing that dynamic changes in chromatin organization could influence viral expression and persistence (40). Furthermore, genome organization assessment in patient-derived CD4^+^ T cells suggested that distinct chromatin accessibility adjacent to intact proviruses could contribute to a selective advantage for the virus during long-term ART (41). Indeed, recurrent genic integration into highly accessible/expressed genes, including several with the capacity to facilitate clonal expansion of the infected cells, has been identified in blood cells from patients on ART (19, 21, 22, 31, 42). Collectively, these studies in blood cells highlight an intimate link between chromatin organization and HIV-1 transcription control, suggesting that new epigenetic targets could serve as candidates for HIV-1 cure research.

While there is currently no precise understanding of the nature of HIV-1 integrations in the brain of PLWH, recent studies in HIV-negative patients mapped regulatory elements in major cell populations of the brain, highlighting the role of enhancers and the 3D genome in neurological diseases (43, 44). This lays ground for an in-depth genomic analysis of the HIV-1 integration profiles in microglia.

Here we report 4,590 integration sites obtained for the first time from HIV-1 infected immortalized microglia cells. We use ChIP-Seq and RNA-Seq to show that proviral integration patterns bearing signatures of genic enhancers found in T cells are corroborated in the microglia background. Chromatin accessibility analysis by ATAC-Seq reveals that CTCF signatures mark latent infection. Furthermore, nuclear architecture exemplified by TAD boundaries fosters HIV-1 insertion into active chromatin. Our findings complement the knowledge of the main HIV-1 CNS reservoir and further highlight the role of dynamic chromatin organization in HIV-1 integration that could contribute to the future therapy development for HIV-1 latent sanctuaries.

## MATERIAL AND METHODS

### Cell Culture and HIV-1 infection

C20 microglia cell line (kindly provided by Dr Alvarez-Carbonell) was cultivated in BrainPhys Neuronal medium supplemented with 10% FBS, 1% penicillin–streptomycin, 500 mg Normocin, 1% N2-Supplement and 1% L-Glutamine at 37°C and 95% CO2.

Approximately 1×10^6^ cells were infected with 250 ng p24/1 million cells of vesicular stomatitis virus glycoprotein (VSV-G) pseudotyped pNL4.3 WT virus through spinoculation for 90 min at 2300rpm and 37°C. Infected cells were cultivated for 3 days at 37°C and 95% CO2.

Viral stocks were produced by transfecting either 25 μg of pNL4.3 viral DNA or 45 μg HIV-GKO (Addgene Plasmid#112234) viral DNA together with 10 μg pMD.G packaging plasmid (45) in HEK 293T cells and collecting supernatants after 72 h following sucrose gradient purification of virus particles. Viral stocks were diluted in PBS, treated with DNAse (15 U/ml, Thermo Fisher Scientific) for 2 h at 37°C, inactivated with 5 mM EDTA before storing them at −80°C.

Productive infection was measured by FACS analysis on BD FACSCelesta™. Therefore, cells were harvested by trypsinization and fixed with 4% PFA for 90 min. After washing with 2% FBS in PBS, HIV-1 p24 was stained with Anti-HIV-1 p24-PE (KC57) (Beckman Coulter) for 30 min on ice and washed again 3 times before measuring on the BD FACSCelesta™.

### Integration assay by *Alu PCR*

The amount of integrated HIV-1 vDNA was quantified through *Alu*-PCR (46). Briefly, in a nested PCR, an integrated virus is amplified with an *Alu* specific primer and a HIV-1 LTR specific primer in the first reaction (Supplementary Table S1). In the second quantitative polymerase chain reaction (qPCR), proviral sequences are amplified using lambda-specific primer (λT) and an internal LTR primer. First round PCR was performed with Alu1 primer (0.1 μM), HIV LTR specific primer LM667 (0.3 μM), 10 mM dNTPs, 10x buffer, AmpliTaq DNA polymerase (Thermo Fisher Scientific) and 100 ng DNA. PCR was performed using SimpliAmp Thermal Cycler (Thermo Fisher Scientific) with the following reaction parameters: 94°C, 15 sec – 94°C 15 sec, 55°C 30 sec, 70°C 2 min – 20x, 72°C 10 min.

The first PCR reaction was diluted (1:50), and the second round was performed by qPCR using iQ™ SuperMix (BioRad), lambda-specific primer λT, internal LTR primer LR and TaqMan probe (Supplementary Table S1). The same qPCR was performed with 10 ng genomic DNA and a housekeeping gene, i.e., lamin B2, B13 region (Supplementary Table S1) used to normalize relative integration levels by the ΔΔC_t_ method (47). Reaction was performed on a CFX96 C1000 Touch Thermal Cycler (BioRad) in triplicates with the following conditions: 98°C 3 min – 98°C 10 sec, 60°C 40 sec – 45x, 98°C 10 min.

### Gene expression analysis by qPCR

RNA was isolated from uninfected and HIV-1 infected cells (WT and HIVGKO) 3 dpi with NucleoSpin RNA Mini kit (MacheryNagel). 500 ng of RNA were reverse transcribed using High-Capacity cDNA Reverse Transcription Kit (Applied Biosystems) according to manufacturer’s instructions. Obtained cDNA was diluted 1:10 and subjected to qPCR analysis in triplicates using iQ™ SuperMix (BioRad). Viral mRNA expression was measured with HIV-1 gag primer probe mix (Supplementary Table 9) and normalized to Eukaryotic 18S rRNA Endogenous Control (VIC™/TAMRA™ probe, primer limited) (Applied Biosystems). Relative expression was calculated using the ΔΔC_t_ method (47). Statistical analysis was performed using GraphPad Prism.

### RNA Sequencing

RNA was isolated from 2.5×10^6^ untreated microglia cells (3 biological replicates) with NucleoSpin RNA Mini kit (MacheryNagel) and sent for RNA sequencing at the Next Generation Sequencing Core Facility of Heidelberg University. Library preparation and rRNA depletion was performed at the facility. Sequencing was performed on a Next Seq instrument, Mid output 2×150bp.

### Chromatin immunoprecipitation sequencing

10×10^6^ million microglia cells were grown in a 15 cm cell culture dish. Medium was removed and cells washed once in PBS (+1 mM sodium butyrate for H3K27ac IP to block deacetylases). Cells were fixed with 1% formaldehyde/PBS (1 mM sodium butyrate for K27ac) for 7 min at RT followed by quenching with 0.125 M glycine/PBS for 7 min at RT. After removing all liquid, 2 times 5 ml ice cold PBS was added to scrape cells and cells were pelleted by centrifugation (1200xg, 7 min). After washing with 10 ml of cold PBS, pellet was resuspended in swelling buffer (10 mM HEPES/KOH pH 7.9, 85 mM KCl, 1mM EDTA, 0.5% IGEPAL CA-630, 1x protease inhibitor cocktail (Roche)) and incubated for 10 min rotating at 4°C. Pellet was dounced 10x before centrifugation 10 min, 3500xg for 10 min at 4°C. One extra wash with swelling buffer without IGEPAL CA-630 was performed before resuspending the nuclei in cold sonication buffer (TE pH=8, 0.1% SDS, protease inhibitor tablet). Sonication was done with a Covaris S220 Focused Ultrasonicator for 18 min (Duty cycle 20%, Intensity 5, Cycles/burst 200). DNA size was followed by agarose gel. DNA fragments should be between 200-500 bp. Triton-X was added to the lysate to a final concentration of 1% and incubated for 10 min on ice. Lysate was cleared by centrifugation at 18000xg 4°C for 5 minutes. Magna ChIP Protein A and G magnetic beads (Millipore) were washed twice with TE 0.1% SDS and 1% TritonX and added to the lysate to preclear for 1h at 4°C rotating. 2-8 mg of chromatin were incubated with corresponding amounts of antibody overnight at 4°C. 1% of chromatin was saved as input. The used antibodies were the following: H3K36me3 (ab9050, Abcam), H3K27ac (ab4729, Abcam), H3K27me3 (C36B11, Cell Signaling), H3K4me1 (ab8895, Abcam), H3K9me3 (ab8898, Abcam), H3K9me2 (ab1220, Abcam), Rabbit control IgG (ab46540, Abcam), Mouse control IgG2a (ab18413, Abcam). The next day, protein A and G magnetic beads were washed twice with a sonication buffer plus 1% TritonX and incubated for 2 h at 4°C with the lysates. Beads were washed twice 10 min with cold buffer I (150 mM NaCl, 1% Triton X-100, 0.1% SDS, 2 mM EDTA, protease inhibitor cocktail (Roche)), once 10 min with cold buffer II (10 mM Tris/HCl pH 7.5, 250 mM LiCl, 1% IGEPAL CA-630, 0.7% Deoxycholate, 1 mM EDTA, protease inhibitor cocktail (Roche)), twice 10 min with cold TET buffer (10 mM Tris/HCl pH7.5, 1 mM EDTA, 0.1% Tween-20, protease inhibitor cocktail (Roche)) and eluted with TE buffer, 1% SDS, 100 mM NaCl. 0.5 mg/ml Proteinase K were added, and samples were incubated for 2 h at 55°C and overnight at 65°C. The next day, 0.33 mg/ml RNAse A (Thermo Fisher Scientific) was added and incubated for 1 h at 37°C. Supernatant was removed from the magnetic beads and DNA was purified with AMpure beads XP clean up according to manufacturer’s instructions. Concentrations were determined by Qubit Fluorometer and enrichment was determined by qPCR on a CFX96 C1000 Touch Thermal Cycler (BioRad) with SsoFast™ EvaGreen® Supermix (BioRad) and the following primers: Human Negative Control Primer Set 2 (ActiveMotif), Human Positive Control Primer Set GAPDH-1 (ActiveMotif), Human Positive Control Primer Set MYT1 (ActiveMotif), Human Positive Control Primer Set ACTB-2 (ActiveMotif), SimpleChIP® Human Sat2 Repeat Element Primers (CellSignaling). ChIP libraries were prepared using NEBNext^®^ Ultra™ II DNA Library Prep Kit for Illumina^®^ (NEB) and NEBNext^®^ Multiplex Oligos for Illumina^®^ (Index Primers Set 1) (NEB) according to manufacturer’s instructions. Libraries were sequenced at c.ATG sequencing core facility at Tübingen University on a NextSeq instrument 2×75 bp.

### Integration site sequencing

Infected C20 microglia cells were harvested at 3 dpi and DNA was isolated with Qiagen blood and tissue kit according to manufacturer’s instructions. DNA was sonicated with Covaris S220 Focused Ultrasonicator 450 s (Duty factor 10%, 200 cycles per burst) and size was monitored by agarose gel analysis.

The LM PCR protocol was adapted from (48). Sonicated gDNA ends were repaired using End-It™ DNA End-Repair Kit (Epicentre) according to manufacturer’s instructions. Sample was purified with PCR purification Kit (MacheryNagel) according to manufacturer’s instructions. A-tailing was performed using NEBNext® dA-Tailing Module (NEB). The reaction was incubated in a thermal cycler for 30 minutes at 37°C and purified with PCR purification Kit (MN) according to manufacturer’s instructions. An asymmetric double stranded linker was annealed overnight at 12°C (800 U T4 ligase, 10x ligase buffer, 1.5 mM linker). After purification with PCR purification kit or 0.9x AMpure XP clean up beads according to manufacturer’s instructions, first nested PCR with linker and LTR specific primers was performed. 25 μl PCR reaction was set up with 1 mM LTR specific primer, 0.2 mM linker specific primer, 5x buffer, 2.5 mM dNTPs, Phusion polymerase (NEB) and 100ng ligation reaction. The thermal cycler program was the following: 94°C 2 min, −94°C 15’, 64°C 30’, 72°C 30’ – 25x, 72°C 10 min. PCR reactions were purified with PCR purification kit (MN) according to manufacturer’s instructions (or 0.9x AMpure XP clean up beads). The second nested PCR with the same linker specific primer and an inner LTR specific primer containing Illumina sequencing index was performed under the same conditions as the first PCR, with increased cycle number (30 cycles). PCR reactions were purified with PCR purification kit (MN) according to manufacturer’s instructions (or 0.9x AMpure XP clean up beads). Quality of the library was analyzed by Bioanalyzer and NEBNext^®^ Library Quant Kit for Illumina^®^ (NEB). IS libraries were sent for sequencing to c.ATG sequencing core facility at Tübingen University and sequenced on a MiSeq instrument 2×150 bp.

### FACS sorting

Approximately 24 million C20 microglia cells were infected with 250ng p24 HIV-GKO virus/ 1 million cells by spinnocculation (2300 rpm, 1.5 h, 37°C). 3 dpi, cells were washed with PBS, trypsinized and filtered in 2% FBS/PBS through a 45 mm filter before sorting on BD FACSAria™ Fusion Cell Sorter into 3 populations: FITC^-^/PE^-^, FITC^+^/PE^+^ and FITC^-^/PE^+^. For each condition, about 80’000 cells were sorted. For the FITC^-^/PE^+^ population 8’000 to 45’000 cells were sorted. 5’000 cells were resorted to check for sorting purity and used for ATAC-Seq as in (49).

### ATAC-Sequencing

ATAC-sequencing was performed as in (49). Cells were pelleted at 500xg, 4°C for 5 min, washed once with cold PBS and resuspended in 50 μl cold lysis buffer (10 mM Tris·Cl, pH 7.4, 10 mM NaCl, 3 mM MgCl_2_, 0.1% (v/v) Igepal CA-630). The reaction was centrifuged immediately for 10 min at 500 × g, 4°C. Supernatant was removed and transposition reaction was set up with 25 μl TD (2x reaction buffer, Nextera Kit, Illumina), 2.5 μl TDE1 (Tn5 transposase, Nextera Kit, Illumina) and 22.5 μl nuclease free H_2_O. The reaction was incubated for 30 min at 37°C shaking on a thermal block (350 rpm). DNA was purified with Zymo ChIP clean and concentrator kit (Zymo research) according to manufacturer’s instructions. Eluted DNA fragments were amplified by PCR with 1.25 μM Primer 1, 1.25 μM Primer 2 (Supplementary Table S1), 25 μl NEBNext® Ultra™ II Q5® Master Mix (NEB) and the whole purified DNA sample (5 min 72°C, 30 sec 98°C – 10 sec 98°C, 30 sec 63°C, 1 min 72°C – 5x). To determine the remaining cycle number for appropriate amplification, 5 μl of the pre-amplified reactions were amplified by qPCR with SsoFast™ EvaGreen® Supermix (BioRad) and 0.25 μl (25 μM) Primer 1, 0.25 μl 25 μM Primer 2 (30 sec 98°C, - 10 sec 98°C, 30 sec 63°C, 1 min 72°C – 20x). Linear RN was plotted versus cycle number to determine the cycle number corresponding to one-third of the maximum fluorescent intensity. The remaining PCR reaction was run with the determined cycle number with the same cycler conditions as before and DNA was purified with Zymo ChIP clean and concentrator kit (Zymo research) according to manufacturer’s instructions. Libraries were sequenced at c.ATG sequencing core facility at Tübingen University on a NovaSeq instrument 2×50 bp.

### Immuno HIV-1 DNA Fluorescent in situ hybridization

Approximately 1.5×10^5^ microglia cells were plated on coverslips on the day of infection in a 24-well plate. 3 dpi cells were washed with PBS and fixed in 4% PFA in PBS for 10 min. Coverslips were extensively washed with PBS and cells were permeabilized in 0.5% triton X-100/PBS for 10 min. Cells were washed three times with PBS-T and incubated in 0.5% triton X-100/0.5% saponin/PBS for 10 min. After three washings with PBS-T, coverslips were treated with 0.1 M HCl for 10 min, washed three times with PBS-T and additionally permeabilized with 0.5% triton X-100/0.5% saponin/PBS for 10 min. After another three washings with PBS-T, RNA digestion was performed with RNAse A (100 μg/ml) for 30 min at 37° C. Coverslips were equilibrated for 5 min in 2x SSC and incubated in hybridization solution (deionized Formamide, 20x SSC, sterile water, pH 7.0) overnight at 4°C. HIV-1 FISH probes were generated by labeling HIV-1 DNA (pHXB2) plasmid. Biotin-dUTP nucleotide mix containing 0.25 mM dATP, 0.25 mM dCTP, 0.25 mM dGTP, 0.17 mM dTTP and 0.08 mM biotin-16-dUTP in H_2_O was prepared. 3 μg of pHXB2 were diluted with H_2_O in a final volume of 12 μl, and 4 μl of each nucleotide mix and Nick translation mix were added. Labeling was performed at 15° C for 5 h. Probes were precipitated in 100% ethanol with sodium acetate, herring sperm and human COT DNA overnight and resuspended in 2x SSC/10% dextran sulfate/50% formamide, denatured and stored at −20° C until use. Probe hybridization was performed with 2 – 4 μl of HIV-1 probe in 6 μl reaction with 2x SSC/10% dextran sulfate/50% formamide, denatured at 95° C for 5 min and then kept on ice for 1 min. After spotting the probe on a glass slide and sealing the coverslip with rubber cement, the slide was put at 80°C for 5 min on a heating plate for denaturation. Subsequently, the hybridization reaction was incubated at 37°C for 48 to 72 h in a closed dish in an incubator. Probe detection was carried out by washing the coverslips with 2x SSC and 0.5x SSC at 37°C and 65°C, respectively; 1hr blocking in TSA blocking buffer (TNB) and detection with streptavidin-HRP in TNB (1:1500) for 40 min at 37° C. Coverslips were then washed with TNT wash buffer four times at RT, before incubation with Fluorescein Plus amplification reagent (1:500 in TSA Plus amplification diluent, part of TSA Plus Fluorescein kit) for 5 min at RT. After 5 final five washings with TNT buffer and nuclear counterstaining using 1:10000 Hoechst 33342 in PBS followed by two washings in PBS, coverslips were mounted with mowiol. For confocal microscopy and manual image analysis, 3D stacks were acquired with a Leica TCS SP8 confocal microscope using a ×63 oil immersion objective.

### siRNA knockdown of LEDGF/p75 and CPSF6

The day before transfection, 2× 10^5^ microglia cells were seeded in 6-well plates to ensure a confluency of 50% on the day of transfection.

Transfection was performed with jetPRIME® (Polyplus transfection®) according to the manufacturer’s instructions. In brief, 110 pmol (for 50 nm siRNA end concentration) of either ON-TARGETplus Non-targeting Control siRNA (Horizon Discovery), ON-TARGETplus Human PSIP1 siRNA SMARTPool or ON-TARGETplus Human CPSF6 SMARTPool (Horizon Discovery) were mixed with 200 μl jetPRIME® buffer. Then 4 μl jetPRIME® reagent were added to the siRNA mix, incubated for 10 min at RT and added dropwise to the cells.

40 h post transfection (or 48 h post transfection for CPSF6 KD), cells were trypsinized, counted and replated for infection. Cell pellets for western blot analysis were harvested. Microglia cells were infected with 250 ng p24/ 1 million of cells of VSV-G pseudotyped HIV-1 WT virus and harvested after 24h of infection for FACS, Western Blot analysis, DNA and RNA extraction.

### MTT assay

Cell viability upon siRNA transfection was measured through the CellTiter 96® Non-Radioactive Cell Proliferation Assay (3-[4,5-dimethylthiazol-2-yl]-2,5 diphenyl tetrazolium bromide (MTT)) (Promega). 1× 10^4^ microglia cells were plated per well in a 96 well plate and transfected with either ON-TARGETplus Non-targeting Control siRNA (Horizon Discovery), ON-TARGETplus Human PSIP1 siRNA SMARTPool or ON-TARGETplus Human CPSF6 SMARTPool (Horizon Discovery) with jetPRIME® (Polyplus transfection®) according to the manufacturer’s instructions for each time point of interest. On the day of transfection and the time point of infection, a new medium plus MTT solution (15 μl) was added to the cells and after 2 h reaction was stopped by the addition of 100 μl of the solubilization/stop solution. Absorbance values at 570 nm were acquired the next day with an Infinite 200 PRO (Tecan, Männedorf, Switzerland) multimode plate reader. Reactions were conducted in triplicate, and the average signal of the triplicates was normalized over the matched untreated controls and expressed as percentage.

### SDS-PAGE and Western Blot analysis

Cell pellets were lysed in 1x Radioimmunoprecipitation assay (RIPA) buffer (Abcam, ab156034) supplemented with 1x protease inhibitor cocktail (Roche) for 10 min at 4 °C and homogenized by sonication in a water bath for 10 min. Protein concentration was assayed using the Micro BCA Protein Assay Kit (Thermo Fisher Scientific) and equal total protein amounts were loaded and run on a precast NuPAGE Bis-Tris 4–12% (Thermo Fisher Scientific) SDS–PAGE at 120 V. Protein transfer was performed with the Trans-Blot® Turbo™ Transfer System (BioRad) using Trans-Blot Turbo Mini 0.2 µm Nitrocellulose Transfer Packs (BioRad). Proteins were transferred on a nitrocellulose membrane for 10 min, 25V, 2.5A. After blocking the membrane for 1 h with 5% milk in 0.1% PBS-Tween (PBS-T) on a rocker at RT, they were incubated with primary antibody overnight at 4°C on a rocker (anti-beta actin (1:10000), a2228, SigmaAldrich; LEDGF/p75 Antibody (1:2000), A300-848A, Bethyl labs, Anti-CPSF6 antibody (1:2000), ab99347, Abcam). The primary antibodies were diluted in 5 ml of 5 % milk in 0.1% PBS-T. The following day, the membranes were washed 3 times for 5 min with PBS-T and incubated with horseradish peroxidase linked secondary antibody (mouse or rabbit) in a 1:5000 dilution in 5% milk in 0.1% PBS-T on a rocker at RT. After washing 3 times for 5 min with PBS-T, proteins were visualized with SuperSignal™ West Pico PLUS Chemiluminescent Substrate (Thermo Fisher Scientific).

### Integration site determination

A BLAT-based pipeline was created to process the LM-PCR raw data. The method was adapted to be used on both single-end (SE) and paired-end (PE) sequencing. Reads with LTR sequence (PE on the first pair/SE) or with Linker sequence (PE, on the second pair) were selected (allowing for 2 mismatches), and trimmed with Cutadapt (v3.2) (50) to improve alignment. Resulting reads shorter than 15bp were excluded. Trimmed reads were converted to fasta format and aligned to a chimeric genome (hg38 and HIV-1 genome) using BLAT (parameters: *-stepSize=6 -minIdentity=97 -maxIntron=0 -minScore=15*) (51).

Only BLAT results that align at least 30 bp (SE) or 10bp (PE) with the genome and where the alignment start was from 0 and the first 5bp were kept (PE/SE). Uniquely mapped reads were kept for further processing steps in both PE and SE. Non-standard chromosomes and internal integrations on the HIV-1 genome were excluded. In the case of multi-mapped reads on SE, only BLAT results where the difference between the longest aligned portion and the second longest were higher or equal to 25bp were kept. On PE, pairs shown to be in the 1 kb vicinity were considered properly paired and kept.

Integrations were considered to be duplicates if the distance in between them was less or equal to 10 bp. PE (N=1,771) and SE (N=2,822) integrations sets were merged (N = 4,590).

Integrations were annotated to the nearest gene using ChIPpeakAnno (v3.24.2) and the GRCh38 annotation package EnsDb.Hsapiens.v86 (v2.99) (52).

Gene ontology analysis (made on genes with genic integrations) was performed using clusterProfiler (v3.18.1) (53).

### ATAC-Seq data processing

Reads were trimmed by TrimGalore (v0.4) with maximum allowed error rate 0.3 and default filtering parameters. Trimmed reads were aligned to the human genome assembly hg38 using Bowtie2 (v2.3) with default settings for paired-end sequencing (54). Duplicated reads were removed. Peaks were called by MACS2 (v2.1) using the *—nomodel* option and false-discovery rate threshold 0.05 on the merged replicates in each condition (55). We kept peaks called by MACS2 with a score of at least 30. Differential peak analysis was performed using the csaw package (v.1.24.3) using a normalization approach based on the enriched genomic regions. Two replicates for each condition were used. Differential regions with an FDR<0.1 were retained. See Supplementary Table S2 for QC values.

Transcription factor footprinting analysis was performed using the TOBIAS toolbox (v0.11.6) with the motifs from transcription factors identified as part of the microglia signature defined by (56, 57). The position weight matrices from the motifs were obtained from the JASPAR database (58).

All profile plots and metagene plots were generated using soGGi (v1.20) from RPKM-normalized bigwigs generated by bamCoverage (59).

CTCF binding dynamics analysis was derived from the TOBIAS toolbox output (*BINDetect* function) binding scores. K-means clustering heatmaps were produced using ComplexHeatmap (v2.6.2) using Spearman distance (over 10 k-means runs) (60). K-means clustering was performed for k ranging from 3 to 7, and the final value selected was 4.

### RNA-Seq data processing

Raw reads were pre-processed with the RNA-Seq nf-core pipeline and aligned to the human genome assembly hg38 (61). TPM values obtained from the stringtie output were transformed into logTPM [log2(TPM/10 + 1)].

Genes were stratified as high expression (top 10% genes with more than 0 logTPM of expression), mid expression (between high and low), low expression (bottom 10% genes with more than 0 logTPM of expression), and non-expressed (all genes with 0 logTPM of expression).

### ChIP-Seq data processing

Reads were trimmed by TrimGalore (v0.4) with maximum allowed error rate 0.3 and default filtering parameters. Trimmed reads were aligned to the human genome assembly hg38 using Bowtie2 (v2.3) with default settings for paired-end sequencing (54). Peaks were called by MACS2 (v2.1) on the merged replicates in each condition using the *—nomodel* option, *broad cut-off=0*.*1*, and false-discovery rate threshold 0.05 (55). See Supplementary Table S2 for QC values.

Super-enhancers were defined using H3K27ac peaks through the findPeaks function from HOMER (v4.10) using the ‘*-style super -o auto*’ parameters (62).

All profile plots and metagene plots were generated using soGGi (v1.20) from RPKM-normalized bigwigs generated by bamCompare (59).

### Random matched controls generation

The set of random matched controls was generated by sampling TSSs obtained from all genes in the annotation package EnsDb.Hsapiens.v86 (v2.99). TSSs sampling was chromosome-dependent to ensure a similar chromosome bias as the real IS set. Two pairs of integrations (up and downstream) from each TSS matching the distance to the nearest TSS on the real IS were generated and then sampled, considering the ratio of genic and intergenic IS in the real set.

### Hotspot and NMF analysis

The hg38 genome was divided into 50 kb long windows using bedtools *makewindows* function (63). The BAM files were then converted to bigWig through bamCompare (on ChIP-Seq datasets) and bamCoverage (on ATAC-Seq and RNA-Seq datasets) using RPKM normalization (59). *multiBigwigSummary* was used to summarize the data over the 50 kb windows (59).

The NMF analysis was performed using the ButchR package (v1.0) over the summarized combined outputs of *multiBigwigSummary* over RNA-Seq, ChIP-Seq, and ATAC-Seq datasets (59, 64, 65). NMF computation was carried out with 10^4^ iterations, 20 initializations, and rank factorization from 2 to 7. Final factorization rank selected, according to observation and quality metrics, was 4. Heatmaps used for the visualization of the H and W matrixes were produced using ComplexHeatmap (v2.6.2) (60).

### ChromHMM analysis

Chromatin 10-state model in microglia was defined using ChromHMM (v1.22) (66, 67). ChIP-Seq (H3K27ac, H3K36me3, H3K4me1, H3K27me3, H3K9me3, and H3K9me2) and ATAC-Seq datasets generated on this work were used as input. BAM files of the input data were binarized using the *BinarizeBam* function in ChromHMM with a 200 bp bin size. The number of states tested (learned with the *LearnModel* function from ChromHMM) ranged from 5 to 15. The final number of states was defined as 10. Heatmaps were produced using ComplexHeatmap (v2.6.2) (60).

### Assessment of Jurkat WT, LKO, IBD, CKD Integration Sites

We used sequencing data from (68) (BioProject accession PRJNA647337, SRA Study identifier SRP273483) for the comparison with TAD boundaries.

Integration sites were mapped from VSV-G pseudotyped HIV-1 luciferase and GFP reporter viruses infected WT Jurkat cells, LEDGF/p75 Knockout (LKO), LEDGF/p75 KO reconstituted with LEDGF/p75 lacking the Integrase Binding Domain (IBD-/-), CPSF6 Knock down (CKD), as well as WT Jurkat cells infected with HIV-1 N74D or A77V capsid (CA) mutants. All code and detailed description of the mapping procedure and downstream analysis can be found at https://github.com/dunjaglavas/IS-mapping. IS were called on human genome build hg19 and lifted over to hg38. Number of IS obtained were the following: WT (N=823,169), LKO (N=65,717), IBD-/- (N=81,346), CKD (N=91,081), N74D and A77V (N=726,694).

### Density analysis along TADs

All density plots represent the density of the number of events (footprints, IS or ChIP-peaks) along the TADs. To map IS to TADs and quantify integration density by region, we collected TADs coordinates obtained from cortex and hippocampus (69) from the 3D Genome Browser (hg38 reference genome) (70). We defined TAD boundaries in these two datasets as the midpoint of the gap region between two consecutive TADs, resulting in 1,446 and 1,481 boundaries in cortex and hippocampus respectively.

To assess the CTCF density, we used CTCF footprints obtained as previously described in uninfected microglia cell lines. To assess the ChIP-Seq density, we considered the narrow peaks called by MACS2. In order to take into account the different lengths of the peaks, we split the peaks into sub-peaks of length 100bp. For CD4^+^ T cells, we used a list of integration sites previously published (21). For TADs, we used a list of consensus boundaries obtained by (38, 69), requiring the boundary to be present in CD4^+^ T cells as well as in two other blood cell types. CTCF binding sites were obtained from ChIP-Seq performed in CD4^+^ T cells (GSE131055) (71). Bootstrap plots were obtained by subsampling 80% of the integration sites.

### Comparison between the TAD boundary distances and genic-related features

IS were extended by 10KB upstream and downstream and partitioned according to *(a)* their overlap with H3K36me3 peaks (2,486 IS overlapping and 2,104 IS non-overlapping) and *(b)* intergenic (N=728) and genic (N=3,862) locations. Distance to the nearest TAD boundary was assessed using the same cortex (N=1,446) and the hippocampus (N=1,481) boundaries called for the density plots.

### TAD alterations assessment

In order to rank TF associated with TAD boundaries, a random forest classification algorithm was applied to transcription factor binding site (TFBS) footprints obtained from ATAC-seq data. Processed BAM files from ENCODE on a set of 5 distinct cell lines (A549, GM12878, HepG2, IMR90, and K562) and 4 tissues (lung, spleen, ovary, and left ventricle) were used to perform TFBS footprinting (72, 73). If existent, BAM for isogenic replicates were merged. Biological replicates were used independently. Validation was done using a randomly selected subset of one tissue (ovary) and two cell lines (HepG2 and K562). Files used are identified in Supplementary Table S3. BAM files were used as input data for the footprinting analysis toolbox TOBIAS (v0.11.6) (57). The position weight matrices from the transcription factor motifs used for footprinting were the ones included on the TOBIAS snakemake example data (N=86 motifs).

The genome was segmented in sliding windows (overlap=1KB) of 5KB length. For each biological sample, windows were scored using the sum of the TFBS predicted to be bound by the corresponding TF. Log-transformed ATAC-Seq counts over the windows were also used as an individual feature. Using this data, a matrix was produced for each of the biological samples and used as input data for model training in the 6 samples selected for training (A549, GM12878, IMR90, lung, ovary, and left ventricle). Classification output variable (*Y* as overlapping a TAD boundary and *N* as not overlapping a TAD boundary) used for training and validation was defined using the boundaries of TADs from the corresponding 9 distinct cell lines or tissues obtained from the 3D Genome Browser (70). A window was classified as part of the positive class Y if it overlapped the midpoint +/-2.5KB between 2 consecutive TADs. For training, all individual matrices from the biological samples were combined into one (28,885,617 rows and 87 predictors).

Model training was performed using *caret* package (tune length = 3 (final mtry was 2), 500 trees, with 10-fold cross-validation) (74). Downsampling was used to correct class imbalance during training. Validation was independently performed on the remaining 3 biological samples (spleen, HepG2, K562). TFs were ranked according to the feature importance (Fig. 4E).

### Gene activity matrix

Gene activities were computed by summing up the ATAC-Seq signal (bigwig files) in the three conditions (uninfected, actively infected and latent infected) in the promoter region (2 kb upstream of TSS) using UCSC hg38 gene annotation, for N=22,665 genes. ATAC-Seq signal was normalized to RPKM using bamCoverage as described before (59). In order to correct for potential bias due to different ATAC-Seq quality between the three conditions, quantile normalization was applied across each condition.

## RESULTS

### Microglia HIV-1 IS map to introns of highly transcribed genes recurrently targeted in T cells

To obtain HIV-1 integration sites in microglia, we used immortalized C20 cells derived from a brain surgery of an adult patient (75). Three days post infection (3 dpi) with VSV-G-pseudotyped HIV-1, HIV-1 integration, mRNA expression and protein production were assayed to confirm productive infection (Supplementary Figure S1A-C). Virus-genome junctions were amplified and sequenced by linker-mediated (LM) PCR sequencing and 4,590 unique IS were obtained using a dedicated processing pipeline (see Methods; Supplementary Table S4). To evaluate genomic characteristics of the integration sites retrieved, we compared microglia integration profiles with published profiles in CD4^+^ T cells (N=13,544) (including our previous study, GEO: GSE134382) and monocyte-derived macrophages (MDM) (N=987) (21, 27). Overall chromosomal distribution of the HIV-1 integrations was comparable between the three cell types and the majority of IS were enriched within introns (58%), with 14.5% being in distal intergenic regions (Figure 1A,B and Supplementary Table S4). Genic integrations displayed higher similarity among T cells and microglia (Jaccard index 0.209) compared to MDM (Jaccard index 0.096) (Figure 1C), with similar gene ontology (GO) terms mostly associated with chromatin modification related processes (Supplementary Figure S1E,F). In addition, highly targeted genes (harboring ≥5 HIV-1 insertions) in microglia were also frequently targeted in CD4^+^ T cells (56.8%) (Figure 1D) and transcriptionally active, as assessed by RNA-Seq (gene stratification described in Material and Methods). Further global transcriptome analysis of microglia targets showed that most integrations (91%) were within highly and medium transcribed genes (Figure 1E, Supplementary Figure S1D). These results suggest that the genic and gene expression requirements for HIV-1 integration in microglia are comparable to major HIV-1 cellular reservoirs (21, 27).

**Figure 1.**
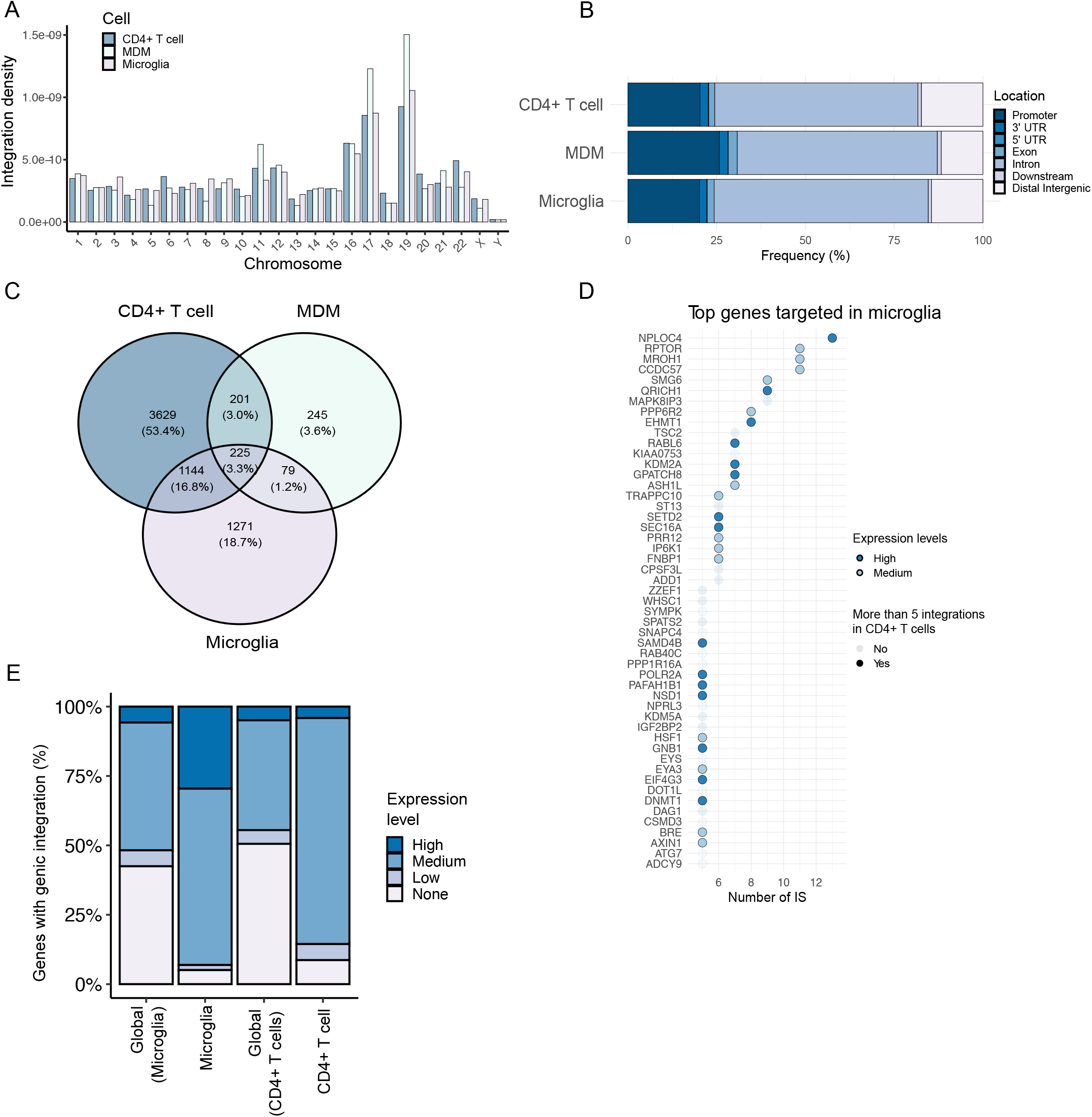
Microglia HIV-1 IS map to introns of highly transcribed genes recurrently targeted in T cells. A) Normalized chromosomal distribution of the integration events for CD4^+^ T cells, MDM, and microglia. Integration density is calculated for each cell type as the number of integrations per chromosome per total number of integrations divided by chromosome length. B) Bar plot showing percentage of IS falling into promoters (from transcription start site (TSS) to 2 kb upstream), 3’ UTR, 5’ UTR, Exons, Introns, Downstream (from transcription end site (TES) to 3 kb downstream) and distal intergenic regions. IS distribution is shown for CD4^+^ T cells, MDM and microglia. C) Shared integrations in CD4^+^ T cells, MDM and microglia. Comparison of the genes between CD4^+^ T cells (N=5199), MDM (N=750), and microglia (N=2719)with at least one integration in their gene bodies. D) Genes with 5 or more integration sites in microglia. Genes were sorted by the number of integration sites found in their gene body. Expression level is highlighted in dark blue (high expressed) and light blue (mid expressed). Fully colored dots denote genes with 5 or more IS in CD4^+^ T cells. E) Bar plot representing the percentage of genic IS according to their expression levels determined by RNA-Seq into high, mid, low or none expressed genes in microglia and CD4^+^ T cells.

### HIV-1 IS in microglia follow H3K36me3 integration signature proximal to genic enhancers

To identify chromatin features of HIV-1 targets in microglia, we mapped histone profiles that accompany gene bodies of actively transcribed genes (H3K36me3), poised/active enhancers (H3K4me1 and H3K27ac), repressive transcription (H3K27me3, H3K9me3), and facultative heterochromatin (H3K9me2) using ChIP-Seq and chromatin accessibility by ATAC-Seq. H3K36me3, bound by the *bona fide* host IN binding protein LEDGF/p75, was enriched at HIV-1 targeted gene bodies with high and intermediate (mid) expression levels, as expected (Figure 2A upper panel). In contrast, the repressive transcription marks H3K27me3 and H3K9me3 were depleted (Figure 2B middle and lower panels), while H3K9me2 signatures remained unchanged (Figure 2B upper panel). Further comparison of genes with (IS) and without integrations (no IS), showed respectively increased trends of H3K36me3 and H3K27me3 occupancy throughout the gene bodies, thus highlighting the opposing role of the two chromatin states in HIV-1 integration (Supplementary Figure S2A).

**Figure 2.**
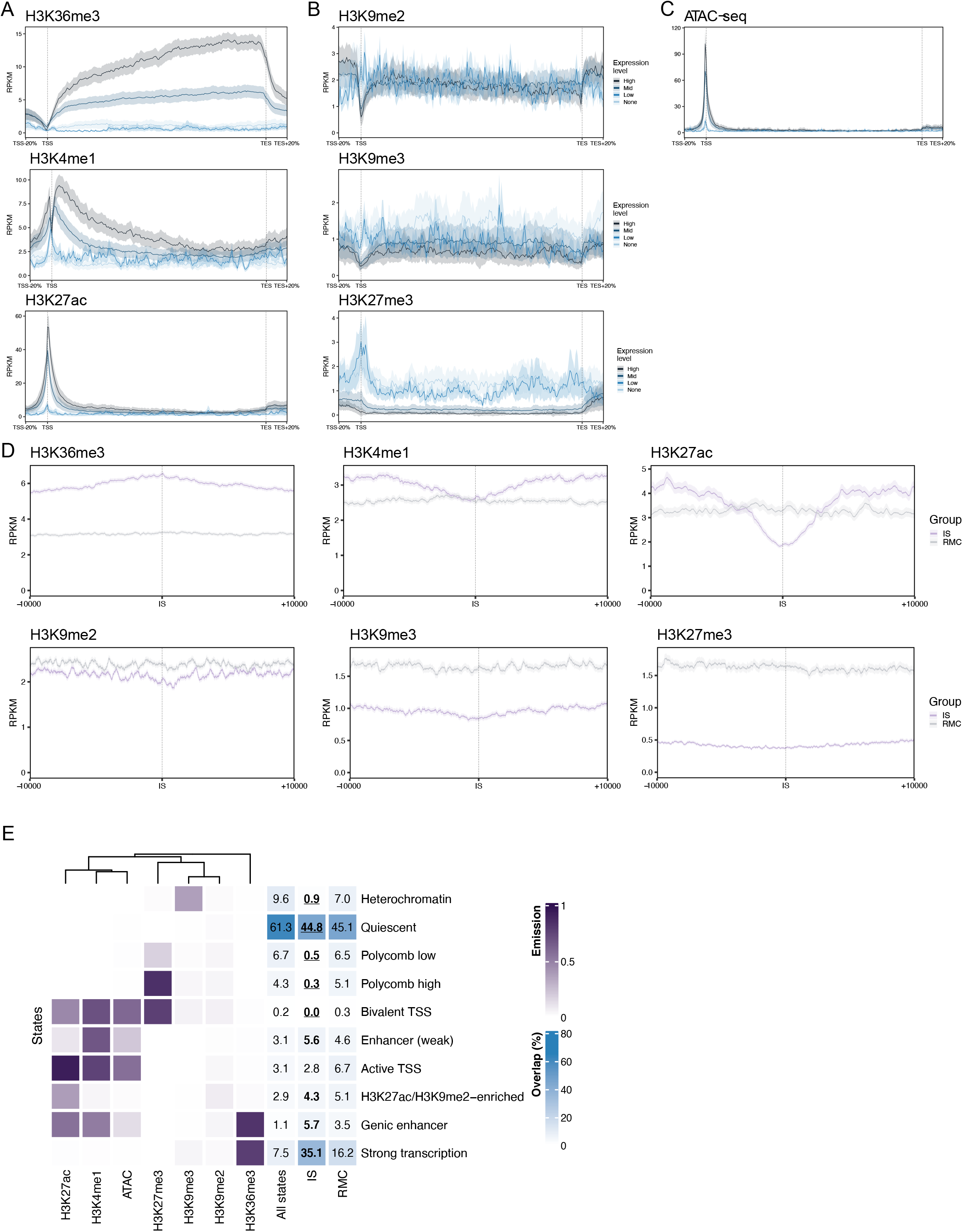
HIV-1 IS in microglia follow H3K36me3 integration signature proximal to genic enhancers. Metagene plots of ChIP-Seq data (in Reads per kilobase per million mapped reads (RPKM)) in genes harboring IS in microglia separated according to their transcription levels into high (black), mid (dark blue), low (light blue) and non expressed (grey). Plots represent the full gene body and 20% of its length upstream of the TSS and downstream of the TES. Confidence intervals (95%) are displayed as the shaded regions. A) H3K36me3, H3K4me1, H3K27ac B) H3K9me2, H3K9me3 and H3K27me3 C) ATAC-Seq signals D) Histone mark profile plots on ChIP-Seq signal (in RPKM) for chromatin marks H3K27ac, H3K4me1, H3K36me3, H3K27me3, H3K9me2, and H3K9me3. Plots represent the integration site vicinity (±10 kb). Microglia integration sites (IS) are shown in purple. Random matched controls (RMC) with the same distance to the nearest TSS and the same intergenic/genic ratio are shown in grey. Confidence intervals (95%) are displayed as the shaded regions. E) Heatmap for emissions (purple) and percentage of overlap (blue) for the chromatin 10-state model generated on ChromHMM for microglia. Each row corresponds to a different state annotated on the basis of the observed results for the modifications and the chromatin accessibility (Left panel). Columns indicate the chromatin features used to generate the model. Higher probabilities for each feature are represented in a darker color (Right panel). Columns indicate the ratios of the states for the entire genome (All states), and the overlap percentage between each chromatin state and the IS (IS) and the RMC set (RMC). Darker colors indicate a larger overlap between the regions represented and the corresponding state. Bold text indicates that the state is targeted by integrations significantly more than expected, bold and underlined text indicates the state is targeted by integrations significantly less than expected (Exact binomial one-sided test, p ≤ 0.05).

Proximal to transcription start sites (TSS) of HIV-1 target genes, H3K4me1 and H3K27ac were enriched at mid and high expression levels, respectively (Figure 2A middle and lower panel). In line with this, ATAC-Seq profiles displayed a highly accessible pattern at TSS, scalable with respect to transcriptional levels (Figure 2C). Furthermore, at TSS of low and not expressed genes, HIV-1 targets were enriched in open chromatin signatures compared to non-targets (Supplementary Figure S2B,C), suggesting that, in addition to transcriptional output, an accessible chromatin state contributes to HIV-1 insertion profiles.

We next assessed chromatin signatures specifically at HIV-1 IS and compared these with randomly generated sites within a +/-10 kb window. H3K36me3 was highly enriched at IS with respect to random matched controls (RMC) and when comparing genic versus intergenic IS positions (Figure 2D, Supplementary Figure S2D). Enhancer histone marks were higher in the vicinity of IS, although not enriched at exact sites, while all repressive histone marks tested were underrepresented (Figure 2D, Supplementary Figure S2D,E).

Finally, we integrated histone profiles and chromatin accessibility datasets generated in this work to comprehensively characterize the microglia IS landscape at high resolution (200 bp, roughly corresponding to nucleosomal scale) using the 10-state ChromHMM chromatin model (Described in Materials and Methods, datasets in Supplementary Table S4) (66, 67). HIV-1 IS patterns were predominantly associated with four chromatin states: active transcription (35.1%), genic enhancers (5.7%), weak enhancers (5.6%), and a H3K27ac/H3K9me2-enriched state (4.3%) (One-sided binomial test, p.value ≤ 0.05). In turn, heterochromatin (0.9%), Polycomb (high and low) (0.3% and 0.5%), and quiescent states (44.8%) were targeted significantly less than expected when compared with the global state ratios (One-sided binomial test, displayed as *All states*, p.value ≤ 0.05) (Figure 2E). In summary, these data indicate that microglia IS are typically found within active regions that are characterized by genic enhancers or regions of high transcriptional output.

### Microglia HIV-1 integrations partition with gene dense transcriptionally active chromatin signatures and are dependent on LEDGF/p75 and CPSF6 host factors

It is well established that one of the defining features of HIV-1 integration are genic insertion hotspots, displaying a local enrichment in IS, identified both in *in-vitro* studies and in patients (19–30). HIV-1 integration association with genic and distal enhancers, including SE, was previously described in T cells (21, 29, 31). To investigate further the previously observed relation of genic enhancers with IS at nucleosomal resolution in microglia (Figure 2E) and to find other IS-associated genomic signatures on a larger genomic scale, we first employed a NMF-based approach recently developed (64). In this analysis, the genome was divided into 50 kb bins and each bin was characterized by its RNA-Seq, ChIP-Seq, and ATAC-Seq signal. The resulting matrix was decomposed using NMF into an exposure matrix (H) and a signature matrix (W), which share an internal dimension. The columns of the exposure matrix (H) were clustered (Figure 3A) and distinct signatures were identified using this approach, as shown in the exposure (Figure 3A) and signature (W) matrices (Figure 3B). Most microglia integrations distributed among 2 signatures: signature 1 (H3K36me3, high transcription/RNA-Seq with 15.3% of bins containing IS) and signature 4 (H3K27ac, H3K4me1, accessible chromatin/ATAC-Seq with 8.3% of bins containing IS), but were underrepresented in signatures 2 (H3K9me2 and H3K9me3 with 1.3% of bins containing IS) and 3 (H3K27me3 with 1.1% of bins containing IS) (Figure 3B). Comparing the locations of SE with the chromatin signatures obtained in the NMF analysis revealed that signature 4 targeted by HIV-1 was also a SE-rich signature (12.7% in sign. 4, compared to 0.8%, 0.1% and 0.2% in the sign. 1, 2 and 3, respectively) (Figure 3B). In addition, proximity between SE and IS was also observed in microglia, implying that this association might not be specific only to T cells (Supplementary Figure S3A) (21, 29, 31).

**Figure 3.**
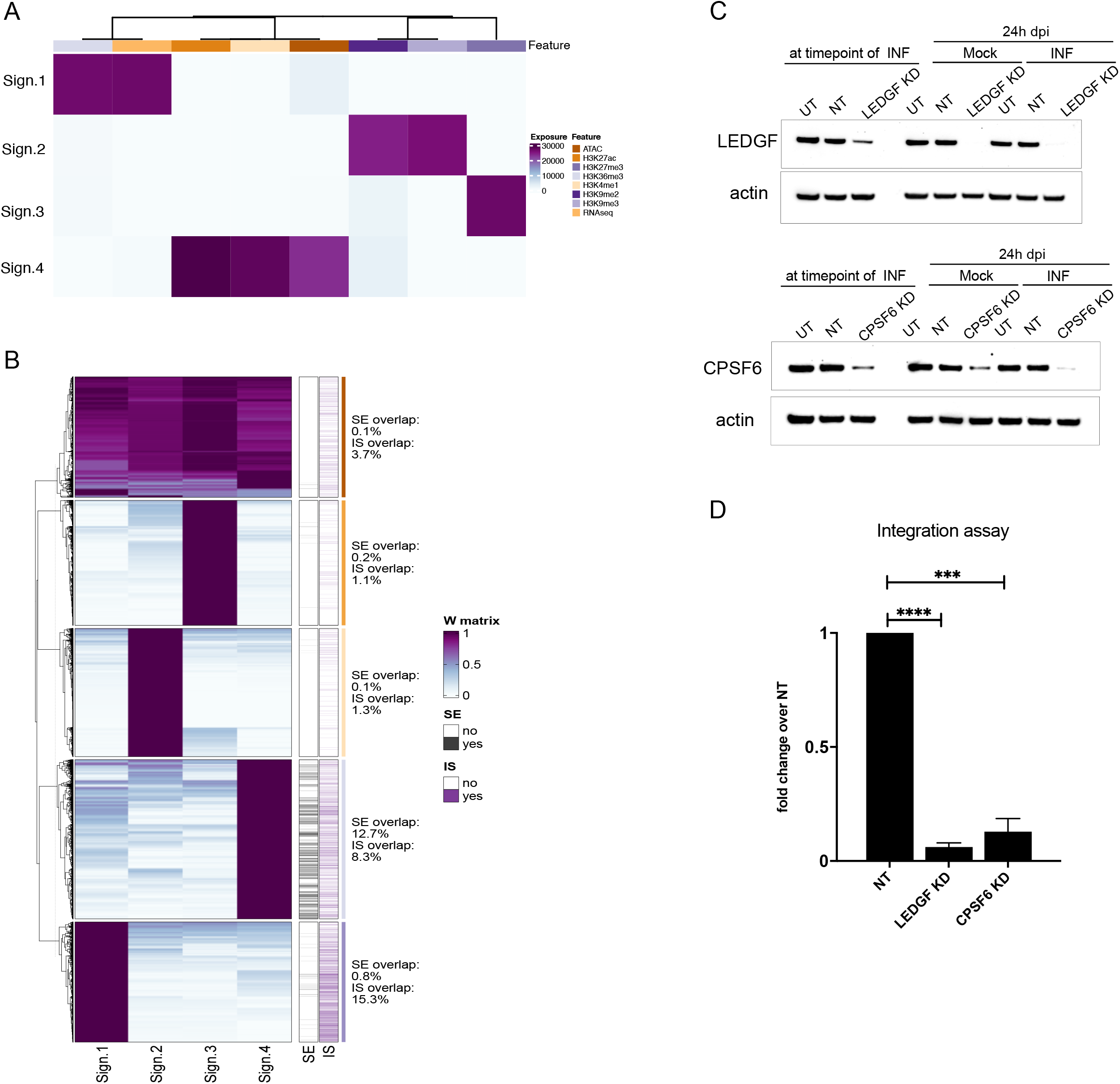
Microglia HIV-1 integrations partition with gene dense transcriptionally active chromatin signatures and are dependent on LEDGF/p75 and CPSF6 host factors. A) Heatmap representation of the H matrix of the NMF performed on the summarized ATAC-Seq, ChIP-Seq, and RNA-Seq signal (8 features) over the 50 kb windows (N=57,238). Colors indicate exposure values of each feature (columns) in the signatures (rows). B) Heatmap representation of the W matrix of the NMF performed on the summarized ATAC-Seq, ChIP-Seq, and RNA-Seq signal over the 50 kb windows (N=57,238). Columns represent the signatures and rows represent the features (50 kb windows). The color indicates the feature contribution to each signature. C) Representative image of western blot analysis of LEDGF/p75 (upper panel) and CPSF6 (lower panel) expression after siRNA knockdown (KD). D) Box plot showing HIV-1 integration levels in microglia cells transfected with LEDGF/p75 or CPSF6 siRNA relative to non-targeting (NT) siRNA control transfection of 3 biological replicates. Data are presented as mean ± SEM and analyzed by unpaired t-test (*** p=0.001, ****p<0.0001).

Two cellular proteins, lens epithelium-derived growth factor (LEDGF)/p75 and cleavage and polyadenylation specificity factor (CPSF) 6, largely determine HIV-1 integration site preferences to gene bodies and transcriptionally active chromatin throughout interaction with viral integrase (IN) and capsid (CA), respectively (76–79). Depletion of either of the integration cofactors diminishes viral integration, with residual insertions mapping away from gene-dense regions of active transcription (31, 79–82).

We thus tested the role of LEDGF/p75 and CPSF6 in the integration process in microglia. Consistent with the enrichment of H3K36me3 at HIV-1 IS (Figure 2A and D), and the overrepresentation of HIV-1 hotspots in the signature enriched in H3K36me3 and with a high transcriptional output (Figure 3B, signature 1), knockdown of either of the proteins led to a significant decrease of HIV-1 integration levels (Figure 3C,D, Supplementary Figure S3B). Hence, HIV-1 integration depends on LEDGF/p75 and CPSF6 in this target cell type as well.

### CTCF marks HIV-1 infection in microglia

To investigate chromatin organization upon HIV-1 infection, we infected microglia with the HIVGKO reporter virus, allowing us to distinguish uninfected cells (eGFP^-^mKO2^-^) from actively (eGFP^+^mKO2^+^) or latently infected cells (eGFP^-^mKO2^+^) using Fluorescence-activated cell sorting (FACS) (Supplementary Figure S4A-E) (83). We then applied ATAC-Seq on the three cell populations (Supplementary Figure S4F) and obtained 92,812/95,401/54,744 peaks in the uninfected, actively infected and latent infected conditions (MACS2 Q-value < 0.001). Global assessment of gene-associated peaks revealed that all three cell populations were characterized by neuronal- and glia-specific biological functions (Supplementary Table S5). Differential analysis of ATAC-Seq signals across the cellular genome displayed modest differences between the three cell populations (Supplementary Table S6). The overall chromatin accessibility remained unchanged at 3 dpi, as only 6, 161 and 379 regions were differentially accessible between uninfected and latent, active and latent and uninfected and active populations, respectively (Supplementary Figure S4G, red dots represent significantly changed regions, false discovery rate (FDR)<0.1). However, using ATAC-Seq based gene activity as a proxy for gene expression, we observed that latent infection was distinguished from actively infected cells by underrepresentation of biological processes involved in cell growth, such as reduced rRNA biogenesis and membrane signaling, and by overrepresentation of neuronal and potassium channel genes (Supplementary Figure S5A and 5B middle column, Supplementary Table S7). In addition, terms related to chromatin modifying enzymes and chromatin organization appeared to be depleted in genes that were highly active in latent versus uninfected cells (Supplementary Figure S5B right column).

Next, we performed differential TF footprinting analysis in the three populations using a recently established framework (57). Our analysis for the enriched TFBS over open chromatin regions (OCRs) identified differential binding of several TF, including HIV-1 transcription factors: nuclear factor ‘kappa-light-chain-enhancer’ of activated B-cells (NFKB), signal transducer and activator of transcription 1 (STAT1) and Activator protein 1 (AP-1) component FOS (colored dots indicate overrepresented footprints in the three states, FDR<0.05, differential binding score >|0.05|) (Figure 4A-C, Supplementary Table S8) (56, 84, 85). NF-KB subunit displayed increased footprints in productive and latent infection compared to uninfected cells, while STAT1 binding was tracked in active versus latent condition (Figure 4A,B). Interestingly, CTCF, an architectural factor involved in 3D genome organization, had decreased TFBS footprints upon productive microglia infection, and higher occupancy in latently infected microglia when compared to productive infection, in line with a recent study showing evidence for CTCF contribution to HIV-1 latency in T cells (40) (Figure 4A-C). We further focused on differentially occupied CTCF binding sites in the three cell populations, and confirmed an overall reduction of CTCF occupancy in productive infection (Supplementary Figure S5C). By selecting only CTCF binding sites occupied in at least one condition, the dynamics of individual TFBS was followed throughout infection (Figure 4D). While some clusters of CTCF binding sites were stable regardless of the infection state (cluster 2-4), others are shown to be more dynamic, displaying on and off states (cluster 1). Of note, stable clusters contained more HIV-1 hotspots (cluster 2 with 2.5%, cluster 3 with 4.4%, cluster 4 with 2.4%, compared to 2.1% of bins containing IS), suggesting a link between HIV-1 integration and CTCF binding. We hypothesized that the role of CTCF in defining TAD boundaries might influence the HIV-1 integration landscape. To test this possibility, we made use of available ATAC-Seq data and corresponding Hi-C datasets to train a Random Forest classification model in order to first curate TF footprints from OCRs and then identify TFs associated with Hi-C derived TAD boundaries (Supplementary Figure S5D). By comparing top-ranked boundary-associated TFs with their global TFBS occupancy in the three microglia cell populations (uninfected, actively and latently infected), we observed that 20 TFs, most prominently CTCF, interferon regulatory factor 1 (IRF1), RE1-Silencing Transcription factor (REST), and Zinc Finger E-Box Binding Homeobox 1 (ZEB1) (>10% predicted importance for the distinction between TAD boundaries and TADs), had differential footprints in the three cell populations (Figure 4E, Supplementary Table S9). CTCF, REST and ZEB1 marked the latent population (FC(latent/active)>0) (Figure 4A,B and Figure 5E), while IRF1 displayed higher enrichment in active cells (FC(latent/active)<0), suggesting that they could potentially contribute to 3D genome organization in the infected cell populations.

**Figure 4.**
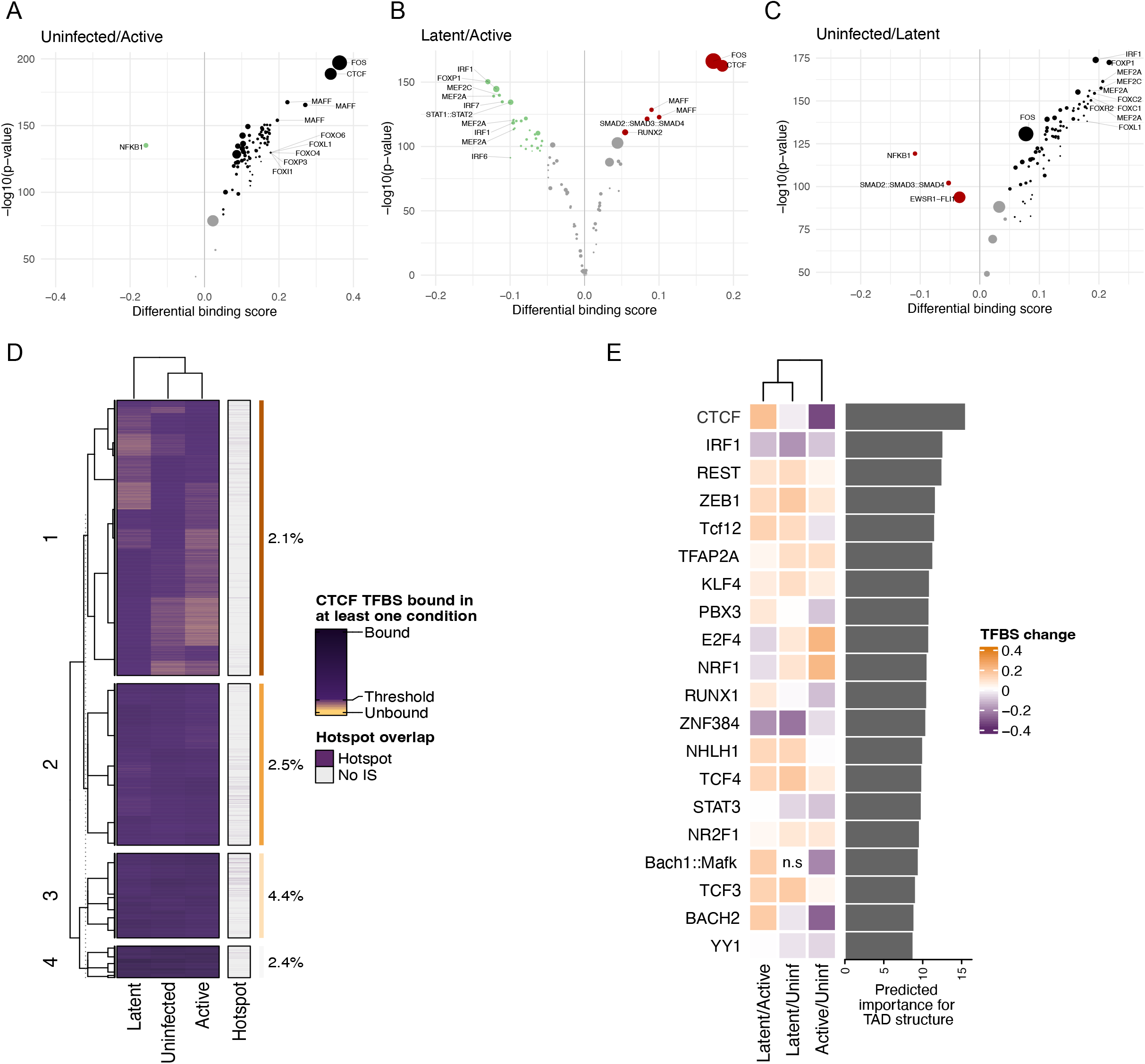
CTCF marks HIV-1 infection in microglia. Volcano plots showing differential footprint accessibility between cell populations. Positive x-values indicate higher accessibility, while negative values indicate lower accessibility in the first mentioned population. Dot size indicates the number of binding sites. A) uninfected/active infection B) latent/active infection C) uninfected/latent infection D) Heatmap displaying the CTCF TF binding dynamics between conditions. Rows represent all CTCF binding sites found to be bound in at least one condition (N=21,727) and columns represent the three conditions. Colors indicate the binding scores (log+1 transformed) obtained from footprinting analysis. The threshold of binding on the legend indicates the score upon which CTCF is considered to be bound. The row clustering is based on the binding scores and it was performed with k-means = 4. E) Heatmap of binding changes for the top 20 TF features (excluding ATAC, observed to be the top feature) of the Random Forest (RF) model. Rows (TF) are ordered by their importance into classifying windows as TAD boundaries and non-TAD boundaries on the training stage of the RF model. Feature importance of each TF feature is shown as a barplot on the right. Colors on the heatmap indicate the degree of TF binding change, compared between the conditions in the columns (orange colors indicate the TF is more bound, whereas purple indicates the TF is less bound for the comparisons shown on the columns).

**Figure 5.**
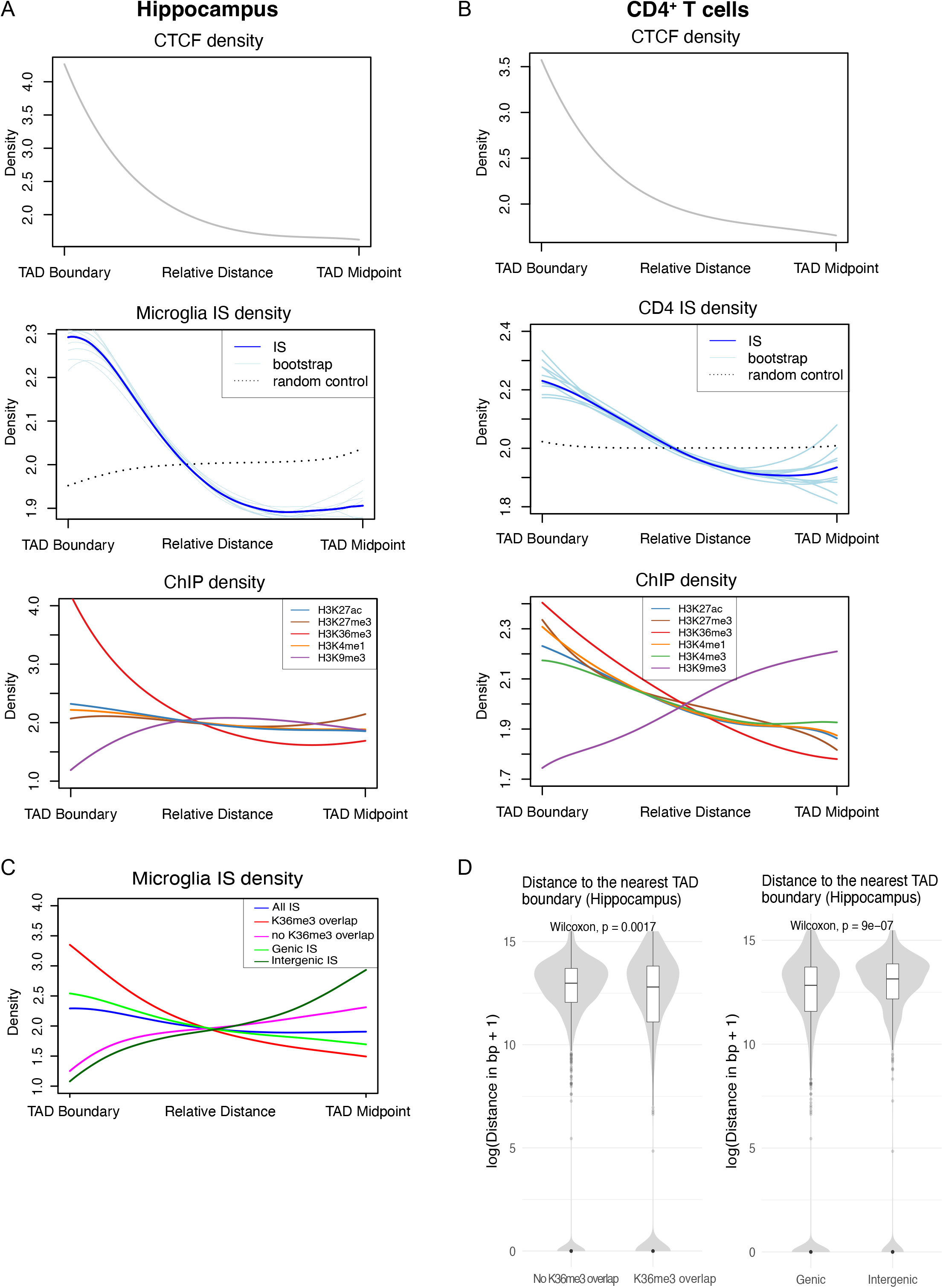
HIV-1 integrates proximal to TAD boundaries. A) Density location plots of microglia CTCF TFBS footprintings (grey line, upper panel), IS in microglia (dark blue line, middle panel), and microglia ChIP-Seq peaks for H3K27me3, H3K36me3, H3K4me1, H3K9me3, and H3K27ac (lower panel) over the boundary-to-midpoint regions of hippocampus TAD boundaries (N=1,481). IS density is compared with random controls (black dotted line) and 10 bootstrapped subsamples (light blue lines) obtained based on the real IS set. B) Density location plots of CD4^+^ T cell CTCF binding sites (grey line, upper panel), IS in CD4^+^ T cells (dark blue line, middle panel), and CD4^+^ T cell ChIP-Seq peaks for H3K27ac, H3K27me3, H3K36me3, H3K4me1, H3K4me3, and H3K9me3 (lower panel) over the boundary-to-midpoint regions of consensus TAD boundaries. IS density is compared with random controls (black dotted line) and 10 bootstrapped subsamples (light blue lines) obtained based on the real IS set. C) Density location plot of microglia IS overlapping (red line, N=2,486) and non-overlapping (pink line, N=2,104) H3K36me3 peaks or intergenic (dark green line, N=728) and genic (light green line, N=3,862) IS over the boundary-to-midpoint regions of hippocampus TAD boundaries (N=1,481). D) Comparison of the distances (log-transformed + 1) from IS overlapping (N=2,486) and non-overlapping (N=2,104) H3K36me3 peaks or intergenic (N=728) and genic (N=3,862) to the nearest hippocampus TAD boundary.

In summary, our comparative ATAC-Seq analysis showed that global chromatin accessibility remained largely unchanged between infection states while the predicted differential CTCF binding, together with other TFs, was identified as a plausible contributor to the 3D genome organization during HIV-1 infection in microglia.

### HIV-1 integrates proximal to TAD boundaries

To further investigate the association of CTCF and 3D genome organization in microglia HIV-1 integration, we used TAD coordinates from published brain tissue Hi-C datasets (hippocampus: GSM2322543 and cortex: GSM2322542 (69)). We first mapped microglia ATAC-Seq derived CTCF footprints and HIV-1 IS from this study to the brain tissue-derived TAD coordinates, using relative distances for each TAD (boundary to midpoint) (Figure 5A, Supplementary Figure S6A). The distribution density plots of CTCF binding sites with respect to hippocampus (TAD boundaries N=1,481) and cortex (TAD boundaries N=1,446) showed a characteristic increase of CTCF binding at the borders (Figure 5A, Supplementary Figure S6A, upper panel). Remarkably, HIV-1 IS distribution (dark blue line) was prominently enriched towards TAD borders when comparing 10 IS subsampling experiments (light blue lines) to the random control sites (black dotted line) (Figure 5A, Supplementary Figure S6A, middle panel). Furthermore, projection of microglia ChIP-Seq profiles generated in this study showed strong enrichment of the gene body and HIV-1 IS-associated histone mark, H3K36me3, in proximity to TAD borders. Densities of genic and active enhancers (H3K4me1 and H3K27ac) and heterochromatin mark H3K27me3 were unchanged with respect to the relative distances within the TAD, while H3K9me3 signal was depleted at the boundaries (Figure 5A, Supplementary Figure S6A, lower panel).

We then asked whether the association between TAD boundaries and HIV-1 IS observed in microglia is maintained in the major blood target cell type, by using TAD coordinates from CD4^+^ T cells together with the integration site collection from the *ex vivo* infections and patient samples we previously curated (Figure 5B, Supplementary Table S4) (86, 21, 38, 71). TAD border regions were confirmed through the mapping of CD4^+^ T cells-derived CTCF ChIP-Seq peaks, as expected (Figure 5B, upper panel, GEO:SE131055 (21, 71)). Notably, enrichment of IS density profiles towards the TAD boundaries was also detected in T cells (dark blue line, Figure 5B, middle panel) (21, 71). Furthermore, densities of H3K36me3, H3K4me1 and H3K27ac chromatin profiles peaked at the boundaries mirroring the IS profile together with active TSS chromatin mark H3K4me3 (Figure 5B, lower panel). Given the strong enrichment of H3K36me3 at TAD boundaries, and the role of IN-LEDGF/p75 interactions with this mark, we analyzed the distribution of IS stratifying by presence/absence of H3K36me3 and genic/intergenic location. Microglia IS not overlapping H3K36me3 peaks (2140 out of 4590 IS) and present at intergenic locations (728 out of 4590 IS) were depleted at TAD boundaries and enriched towards TAD midpoints both on hippocampus and cortex TAD borders (Figure 5C, Supplementary Figure S6B). Moreover, HIV-1 IS proximity to boundaries not overlapping with H3K36me3 or located in intergenic regions was significantly lower when compared to overlapping H3K36me3 peaks or genic integrations, suggesting that observed integration targeting is strongly H3K36me3 dependent (Figure 5D, Supplementary Figure S6C). H3K36me3 driven TAD insertion properties were confirmed in T cell background, by mapping previously sequenced integration sites from LEDGF/p75 knockout (LKO) and IN binding mutant Jurkat cells (IBD), while CPSF6 knockdown (CKD) or mutant viruses with impaired CA-CPSF6 interaction had no effect on the TAD-proximal proviral positioning (68) (Supplementary Figure S6D).

Taken together, these results indicate that the 3D chromatin environment contributes to HIV-1 IS targeting into transcriptionally active chromatin.

## DISCUSSION

In this study we systematically characterized for the first time HIV-1 integration sites and determined transcriptional output and chromatin signatures of HIV-1 targeted genomic regions in the so far unexplored microglia reservoir. We observed high similarities between microglia and blood derived HIV-1 integration patterns with respect to their chromosomal distribution, recurrent genic integration and intronic targeting of highly transcribed genes, suggesting that common HIV-1 site selection regulation and/or transcriptional programs facilitate integration in all HIV-1 targets (Figure 1A-E, Supplementary Figure S1D,E,F) (19–23, 25–27, 29, 31, 68). On a chromatin level, HIV-1 insertions in microglia were accompanied by H3K36me3-marked genes bodies of highly expressed/accessible genes in the vicinity of genic enhancers (marked by H3K4me1 and H3K27ac) and devoid of repressive chromatin, thus confirming previously established integration profiles in blood HIV-1 reservoirs (Figure 2A-D, and Supplementary Figure S2A-D) (19–23, 25–27). Association of HIV-1 integration patterns with enhancers observed in T cells was corroborated in microglia on both nucleosomal and larger genome scale, as genic and SE elements partitioned with signatures enriched for microglia IS (Figure 2E and Figure 3A,B) (21, 77, 87). Integration requirement for the main cellular cofactors of IN and CA (LEDGF/p75 and CPSF6, respectively) is also preserved in microglia infection (Figure 3C,D and Supplementary Figure S3B) (21, 29, 31, 68, 77, 78, 87–90). These general principles of HIV-1 integration established here for the previously unexplored CNS reservoir prompted us to confront our data with the recently curated comprehensive RIGs and recurrently avoided genes (RAGs) lists generated from HIV-1 IS in 10 different cell types (31). Interestingly, microglia target genes shifted towards RAGs, identified earlier to be omitted for viral integration in non-neuronal cells, and displayed neuro-related GO terms for biological processes (Supplementary Table S10, one sixth of RAGs were targeted in microglia). While the observed RAGs targeting in microglia could be attributed to the cell-specific integration pattern, to date our analysis represents the only microglia data set available, and further IS studies need to be conducted in primary microglia to confirm these observations. Although the microglia cellular model used in this study shares the main cellular, chromatin and transcriptional hallmarks of primary microglia, it does not fully recapitulate microglia properties *in vivo* (56, 75). Future experiments would certainly benefit from these valuable samples to confirm CNS insertion patterns which we identified here to be largely preserved in other cell types.

This study also sheds light on the HIV-1 infection and latency processes in microglia. Based on the viral presence and/or gene expression, we identified a number of differentially accessible TF binding sites. In particular, in the productively infected microglia, we found TF with well-established roles in HIV-1 transcription and T cell and MDM activation (NFKB, STAT1) to be more bound, consistent with the overall activated cell state. On the other hand, the host cell nuclear organization factor CTCF marked the latent infection (Supplementary Figure S5C, Figure 4A,B). Stably occupied CTCF binding sites in the uninfected, active and latently infected cell populations were more targeted by HIV-1 compared to the dynamic TFBS (Cluster 2-4 and Cluster 1, respectively), suggesting that CTCF mediated 3D genome organization could support HIV-1 integration and transcription (Figure 4D). More rigorous conclusions regarding 3D dynamics observed by footprinting analysis would merit genome-wide profiling of CTCF protein, which was limited in this study by the low cell number obtained from the infected sorted populations.

We further distinguished active and latent cell populations based on predicted TF importance for the TAD structure (Figure 4E). Factors with a significant contribution to predicted TADs and overrepresented footprints in actively replicating cells, including IRF1 and NRF1, are known to regulate transcriptional programs and chromatin accessibility of innate immune cells and stress responses to viral infection, respectively (91–94). Likewise, TAD structure prediction was also reinforced by TF with enriched footprints in the latent cell population such as transcriptional repressor REST, also known as Neuron-Restrictive Silencer Factor (NRSF), and CTCF that, as expected, had the most prominent TAD contribution score (Figure 4E) (95, 96). To this end, the role of CTCF in viral replication and latency has been suggested for several DNA viruses, and recently for HIV-1 in primary CD4^+^ T cells (40, 97). Because HIV-1 proviruses do not harbour CTCF binding sites, it is plausible that CTCF contributes to viral gene expression through its role as an insulator in TAD border formation and looping interactions between nonadjacent segments of DNA (36, 37). Here we explored this previously overlooked possibility by mapping microglia CTCF footprints and HIV-1 IS onto TAD boundaries from primary neuronal tissues (Figure 5A upper and middle panel, Supplementary Figure S6A upper and middle panel). We found HIV-1 IS in proximity to TAD borders marked by highly enriched H3K36me3 gene body-associated chromatin and depleted of H3K9me3 heterochromatin, characteristic of topological boundary regions, as previously established for transcriptionally active genes (Figure 5A lower panel and Supplementary Figure S6A lower panel) (36, 98). Indeed, proviral insertion enrichment at borders was strongly dependent on genic integration and H3K36me3 gene body histone modification (Figure 5C and Supplementary Figure S6B), while intergenic IS followed H3K9me3 chromatin mark at the TAD margins. Remarkably, we could recapitulate HIV-1 insertion patterns proximal to TAD flanking regions following both gene body and enhancer histone marks by using patient-derived IS and Hi-C and ChIP-Seq datasets from primary CD4^+^ T cells (Figure 5B, middle and lower panel). Furthermore, impaired IN-host factors interaction in T cell background showed loss of TAD border proximal insertion in LKO and IBD cells thus further strengthening an essential role of LEDGF/p75 (and H3K36me3) in recurrent integration targeting (31). Instead, neither CKD nor HIV-1 CA mutants shifted HIV-1 IS away from the TAD boundaries (Supplementary Figure S6D) (31, 68).

Proviral positioning proximal to TAD borders could also mediate the transcriptional outcome upon HIV-1 integration by facilitating ectopic long range contacts. While HIV-1 gene expression is tightly regulated by cellular TFs binding to the 5′ LTR, the chromatin state of the provirus and HIV-1 transactivator protein (Tat)-mediated positive feedback, it also depends on the transcriptional control of promoter activation through the formation of a gene loop joining 5′ LTR promoter and 3′ LTR poly(A) signal (84, 99). In the complex 3D genome organization of a host cell, other looping mechanisms could contribute to the proviral transcription. Enhancer adoption by a newly accessible gene promoter (HIV-1 LTR) upon TAD border disruption could lead to persistent active viral transcription and/or reactivation from latency, a mechanism previously shown to drive oncogenic transformation events (100–102). Alternatively, viral genomes embedded adjacent to TAD boundaries could become silenced by spreading of heterochromatin from neighbouring domains upon interruption of CTCF insulator binding (103, 104). The latter outcome could be also achieved through high transcriptional output at domain borders, which could both contribute to HIV-1 insertion biases toward TAD boundaries and have an insulation effect on HIV-1 transcription leading to latency (36, 39, 98). Either way, the well established metazoan TAD border-enriched active chromatin could support the viral integration into gene bodies as a general feature of HIV-1 integration in different reservoirs (36, 105–107). Although data presented herein collectively show that topological boundaries rather than inner TAD regions represent HIV-1 insertion targets, future studies would benefit from patient derived primary microglia Hi-C to address the role of microglia specific frequently interacting regions in the viral integration and transcription.

Taken together, these results are providing evidence that 3D nuclear architecture complements HIV-1 integration requirements, opening a possibility that TAD border-proximal active chromatin could ensure an appropriate environment for viral replication and/or latency establishment.

## DATA AVAILABILITY

All relevant data supporting the key findings of this study are available within the article and its Supplementary Information files or from the corresponding authors on a request. The integration site, RNA-Seq, ChIP-Seq and ATAC-Seq data will be made available on Gene expression omnibus (GEO) National Center for Biotechnology Information (NCBI) data repository. Scripts used for the computational analysis can be found on https://github.com/hdsu-bioquant/HIVmicroglia.

## SUPPLEMENTARY DATA

Supplementary Data are available at NAR online.

## Authors’ contributions

M.L., B.L., M.R., C.H., and A.L.C designed the research; M.R., B.L., M.K., I.L.S., C.P., and N.T. performed the experiments; M.R., A.L.C., B.L., C.H., D.G. analyzed the data; B.L., M.R., O.T.F., A.L.C., C.H. and M.L. wrote the manuscript; and K.V, O.T.F, C.H., and M.L. provided funding.

We would like to acknowledge Dr David Alvarez-Carbonell, Department of Molecular Biology and Microbiology, Case Western Reserve University, Cleveland, OH, USA for providing microglia cells, Dr Vibor Laketa (DZIF) the Infectious Diseases Imaging Platform (IDIP) platform coordinator for microscopy support, as well as the EMBL Genomics Core Facility Heidelberg and c.ATG Tübingen for their technical support.

## FUNDING

This research was supported by the DFG Priority Program [SPP2202 3D Genome Architecture in Development and Diseases Project “Nuclear landscape of HIV-1 infection in microglia-An unexplored HIV Reservoir” 422856668 to M.L. and C.H.] and by German Center for Infection Research, DZIF [TTU04.820 (HIV reservoir) and TTU04.709 (Preclinical HIV-1 Research) to M.L.] and German Center for Infection Research, DZIF [TTU04.820 (HIV reservoir) to O.T.F.].

DG and KV are supported through the European Structural and Investment Funds grant for the Croatian National Centre of Research Excellence in Personalized Healthcare (contract #KK.01.1.1.01.0010), Croatian National Centre of Research Excellence for Data Science and Advanced Cooperative Systems (contract #KK.01.1.1.01.0009) and Croatian Science Foundation (grants IP-2014-09-6400 and IP-2019-04-5382).

Funding for open access charge: DFG Priority Program [SPP2202 3D Genome Architecture in Development and Diseases Project “Nuclear landscape of HIV-1 infection in microglia-An unexplored HIV Reservoir” 422856668 to M.L. and C.H.]

## CONFLICT OF INTEREST

The authors state no conflict of interests.

## Notes

### Competing Interest Statement

The authors have declared no competing interest.

