## supplementary data file for "Genomic profiling of HIV-1 integration in microglia links viral insertions to TAD organization"

**Supplementary Figure 1.**

3 dpi with HIV-1 VSV-G WT virus cells were harvested for DNA, RNA and p24 protein analysis.

A) Box plot showing one representative replicate of HIV-1 integration levels relative to mock infections, infection inhibited with 10 μM Raltegravir (RAL) and infection inhibited with 20 μM Efavirenz (EFA) (left panel) assayed by Integration assay (*Alu* PCR) (3 technical replicates). Box plot of viral gag mRNA expression relative to mock infection of 3 biological replicates (right panel). Data presented as mean ± SEM.

B) p24 FACS analysis of 3 biological replicates (blue, light and dark orange histograms) of VSV-G WT HIV-1 infections.

C) Three-dimensional immuno-DNA fluorescence in situ hybridization (FISH) of HIV-1 WT in microglia cells (green: HIV-1 probe, blue: DNA staining with Hoechst 33342, scale bar represents 5 μm).

D) Genes with 12 or more integration sites in $CD4^+$ T cells. Genes were sorted by the number of integration sites found in their gene body. Expression level is highlighted in dark grey (high expressed), dark blue (mid expressed), blue (low expressed), and light blue (non-expressed).

E) Dot plot representing the top 15 terms enriched from GO Biological Processes (BP) using the list of genes harboring integrations in microglia.

F) in $CD4^+$ T cells

Rows indicate the enriched GO term and X-axis represents the ratio of genes targeted by HIV-1 in relation to the size of the gene set in the GO term. Dot size indicates the number of genes in each GO term. Color encoding indicates the FDR (≤0.05).

**Supplementary Figure 2.**

Metagene plots of ChIP-Seq signal (in RPKM) on genes harboring integration sites in microglia by gene expression level (N(high)=710; N(mid)=1788; N(low)=64; and N(none)=185). Plots represent the full gene body and 20% of its length upstream of the TSS and downstream of the TES. Dark grey line represents the averaged signal along genes harboring integration sites in microglia. Light blue line represents the averaged signal along a randomly sampled gene set of equal size. Confidence intervals (95%) are displayed as the shaded regions.

A) for chromatin marks H3K36me3 and H3K27me3

B) of ATAC-Seq signal

C) for chromatin marks H3K4me1 and H3K27ac

Histone mark profile plots on averaged ChIP-Seq signal (in RPKM). Plots represent the integration site vicinity (±10 kb) for integrations in microglia and matched random controls subsetted by location as genic (N = 3862, in blue) and intergenic (N = 728, in purple). Genic and intergenic integration sites (IS) are shown in darker blue and purple, respectively. Random matched controls (RMC) with the same distance to the nearest TSS and the same intergenic/genic ratio are shown in light blue (genic) and light purple (intergenic). Confidence intervals (95%) are displayed as the shaded regions for each line.

D) for chromatin marks H3K36me3, H3K4me1 and H3K27ac

E) for chromatin marks H3K9me2, and H3K9me3 and H3K27me3.

**Supplementary Figure 3.**

A) Violin plots displaying the log-transformed distance to the nearest SE for IS and RMC.

B) MTT assay of 3 biological replicates 0 h and 48 h post transfection with either LEDGF/p75 or CPSF6 targeting siRNA normalized to non-targeting (NT) siRNA control.

**Supplementary Figure 4**.

3 dpi with HIV-1 GKO virus, microglia cells were harvested for DNA and RNA analysis.

A) Box plot showing one representative replicate of HIV-1 integration levels relative to mock infections, infection inhibited with 10 µM Raltegravir (RAL) and infection inhibited with 20 µM Efavirenz (EFA) (left panel) assayed by integration assay (*Alu* PCR) (3 technical replicates). Data presented as mean ± SEM.

B) Box plot of HIV-1 viral gag mRNA expression relative to mock infection of 3 biological replicates. Data presented as mean ± SEM.

C) Three-dimensional immuno-DNA FISH of HIV-1 GKO in microglia cells (green: HIV-1 probe, blue: DNA staining with Hoechst 33342, scale bar represents 5 µm).

D) Gating strategy of sorting HIV-1 GKO reporter virus infection. Live cells were gated by forward (FSC-A) and side scatter (SSC-A) analysis. Populations were sorted into uninfected (FITC-, PE-), productive (FITC+, PE+) and latent (FITC-, PE+) infection.

E) Purity control of 5000 sorted cells of uninfected, active and latent populations.

F) Size distribution of mapped ATAC-Seq reads of all replicates.

G) Volcano plots display FC versus FDR of differentially accessible regions of the cellular genome identified in the three cell states. Peaks that are significantly (FDR <0.1) changed are shown in red. Data represent two independent experiments.

**Supplementary Figure 5.**

A) Gene activity was computed by averaging the ATAC-Seq signal over all gene promoters in each of the three conditions; a gene activity matrix and logFC matrix were constructed.

B) Gene set enrichment analysis using Reactome pathways based on the logFC activities between pairwise conditions. Blue/Red dots indicate enrichment/depletion in the first mentioned condition.

C) Venn diagram representing the numbers of CTCF transcription factor binding sites found to be uniquely bound or shared between the three conditions.

D) Schematic workflow of Random Forest Modeling.

**Supplementary Figure 6.**

A) Density location plots of microglia CTCF TFBS footprintings (grey line, upper panel), IS in microglia (dark blue line, middle panel), and microglia ChIP-Seq peaks for H3K27me3, H3K36me3, H3K4me1, H3K9me3, and H3K27ac (lower panel) over the boundary-to-midpoint regions of cortex TAD boundaries (N=1,446). IS density is compared with random controls (black dotted line) and 10 bootstrapped subsamples (light blue lines) obtained based on the real IS set.

B) Density location plot of microglia IS overlapping (red line, N=2,486) and non-overlapping (pink line, N=2,104) H3K36me3 peaks or intergenic (dark green line, N=728) and genic (light green line, N=3,862) IS over the boundary-to-midpoint regions of cortex TAD boundaries (N=1,446).

C) Comparison of the distances (log-transformed + 1) from IS overlapping (N=2,486) and non-overlapping (N=2,104) H3K36me3 peaks or intergenic (N=728) and genic (N=3,862) to the nearest cortex TAD boundary.

D) Density location plots of $CD4^{+}$ T cell IS (red line, N=13,544), Jurkat WT HIV-1 IS (yellow line, N=823,169), HIV-1 N74D and A77V CA mutants IS in Jurkat cells (blue line, N=726,694), IS in Jurkat CKD (green line, N=91,081), IS in Jurkat IBD-/- (violet line, N=81,346), IS in Jurkat LKO (orange line, N=65,717) over the boundary-to-midpoint regions of consensus TAD boundaries.

**Supplementary tables description**

**Supplementary Table S1.** Primer sequences

**Supplementary Table S2.** QC Table for ChIP-Seq and ATAC-Seq

**Supplementary Table S3**. Metadata used for Random Forest Model Training

**Supplementary Table S4.** HIV-1 integration sites in microglia, IS distribution in microglia, MDM and $CD4^{+}$ T cells and data sets used in this study

**Supplementary Table S5.** Gene ontology analysis of ATAC-Seq peaks in uninfected, active and latently infected microglia

**Supplementary Table S6.** Differential ATAC-Seq peaks analysis between uninfected, active and latently infected microglia

**Supplementary Table S7.** Reactome pathway analysis of gene activity matrix

**Supplementary Table S8.** Differential footprinting analysis with TOBIAS of uninfected, active and latently infected microglia

**Supplementary Table S9.** Feature importance obtained in Random Forest Model training for all of the predictors used

**Supplementary Table S10.** GO analysis of microglia integration genes and RAGs

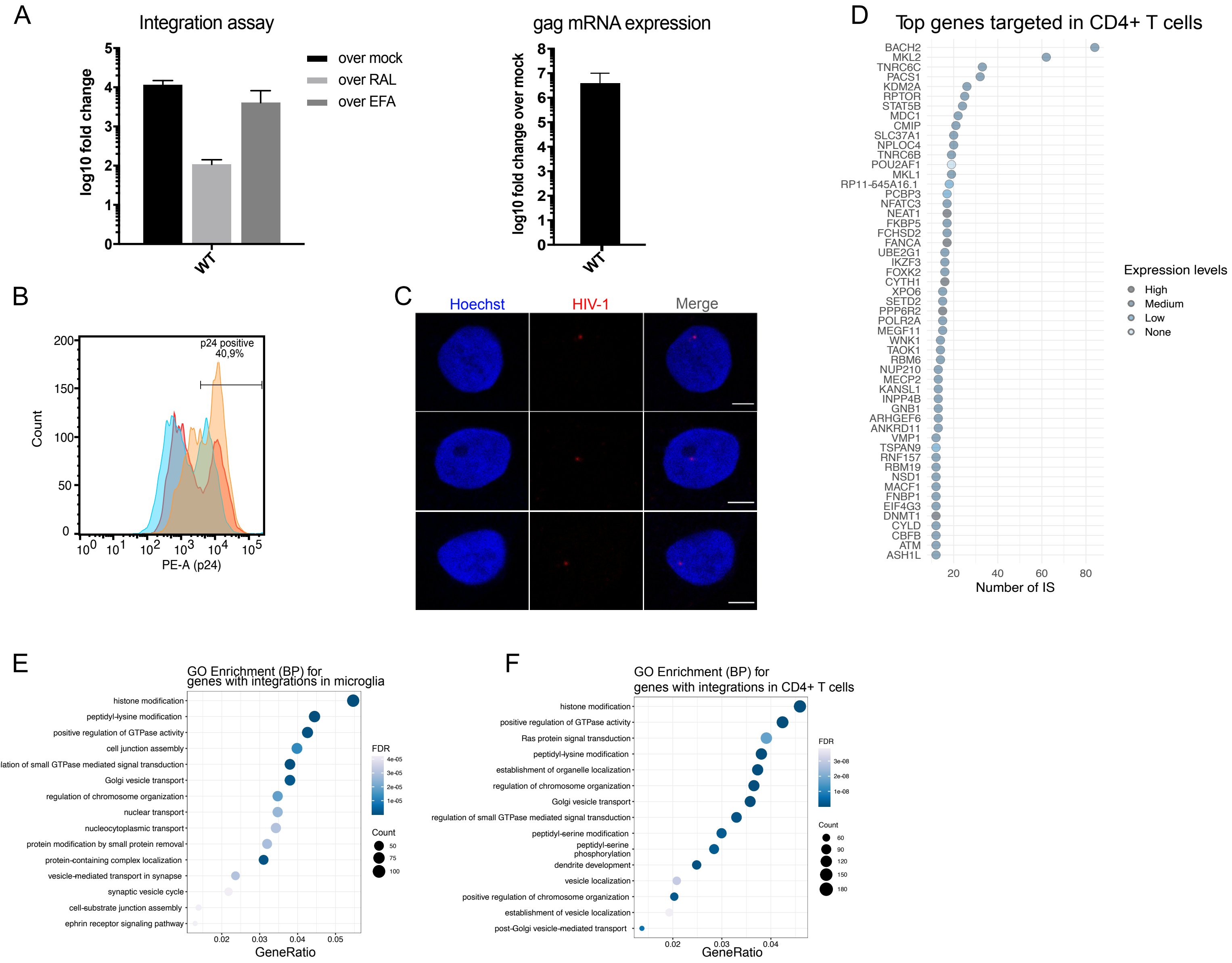


Supplementary Figure S1

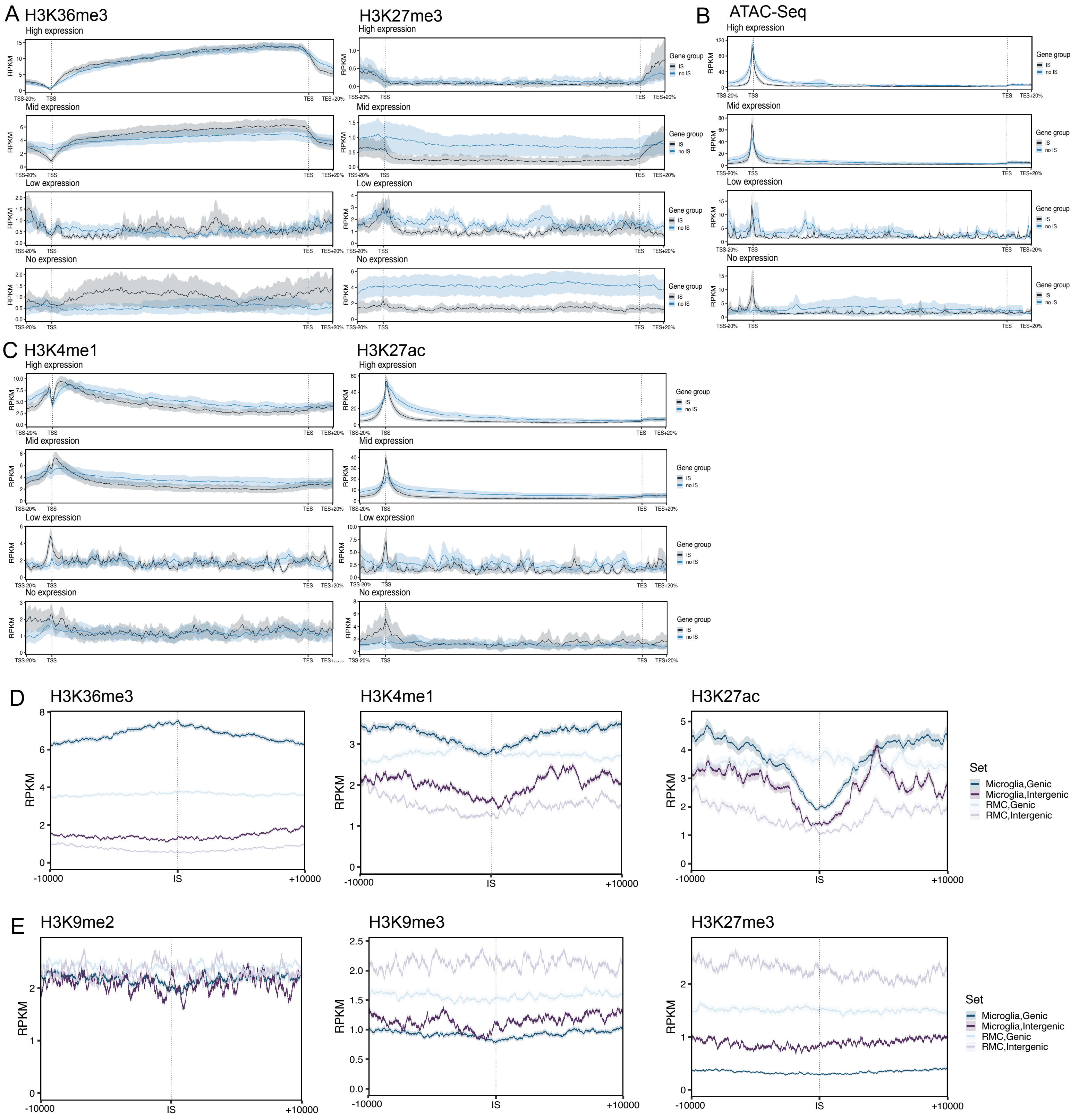


Supplementary Figure S2

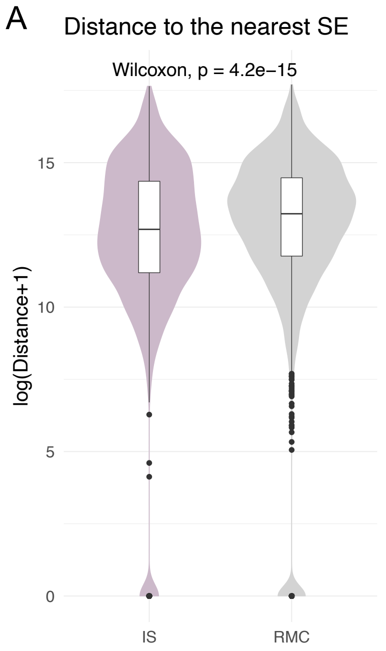


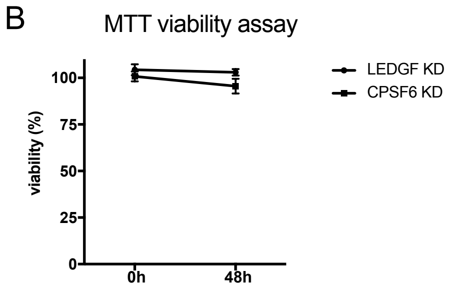


Supplementary Figure S3

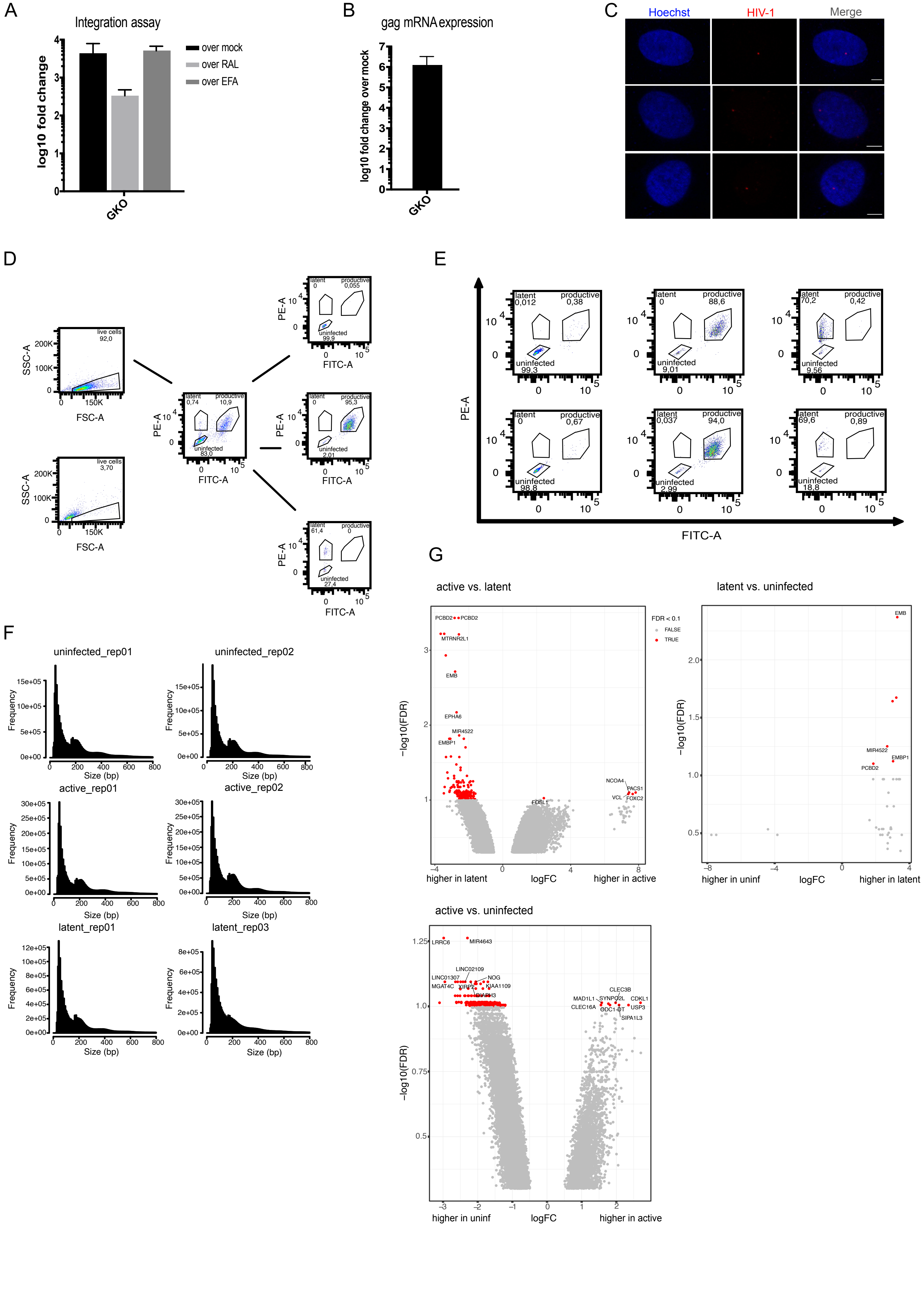


Supplementary Figure S4

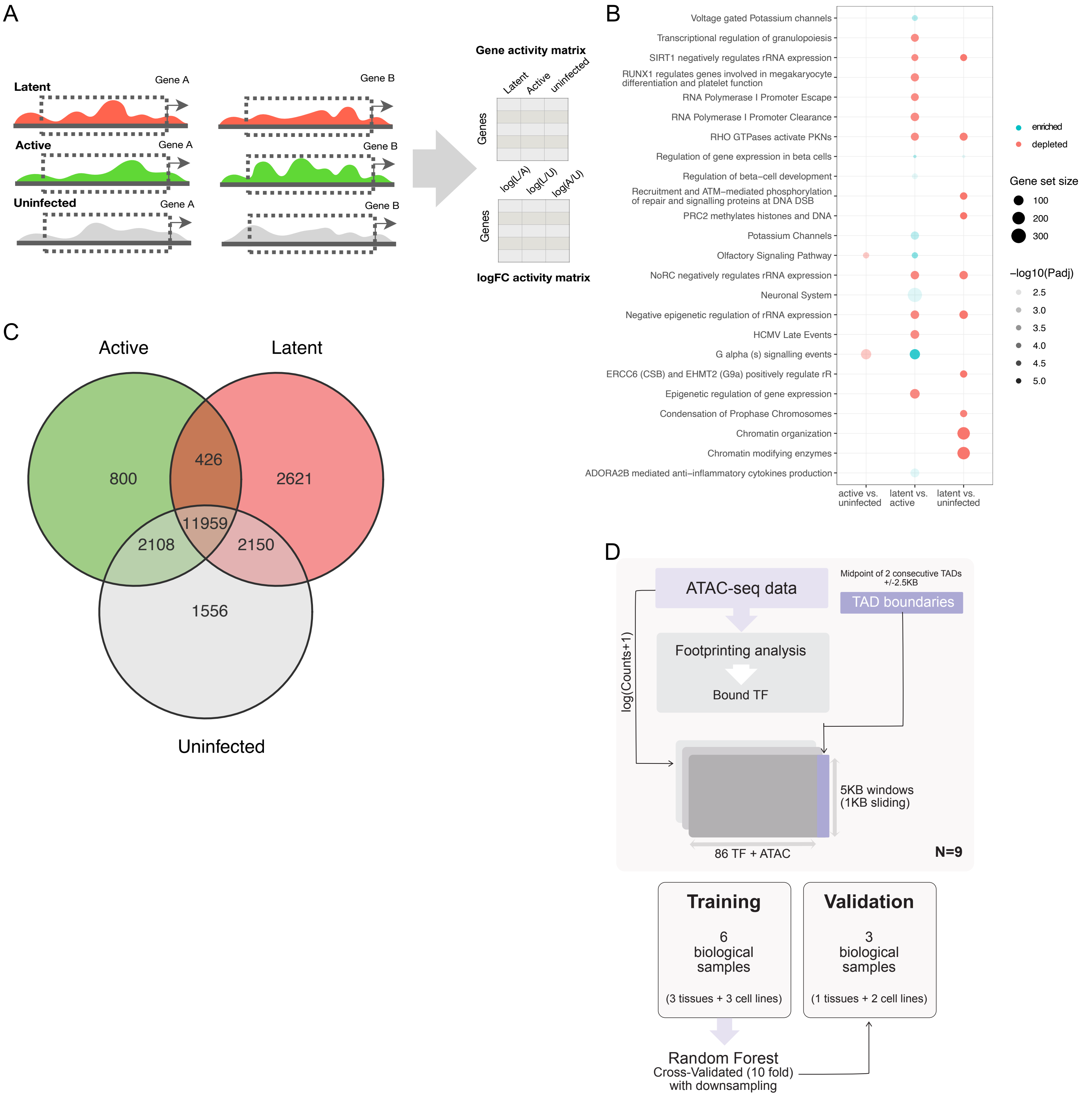


Supplementary Figure S5

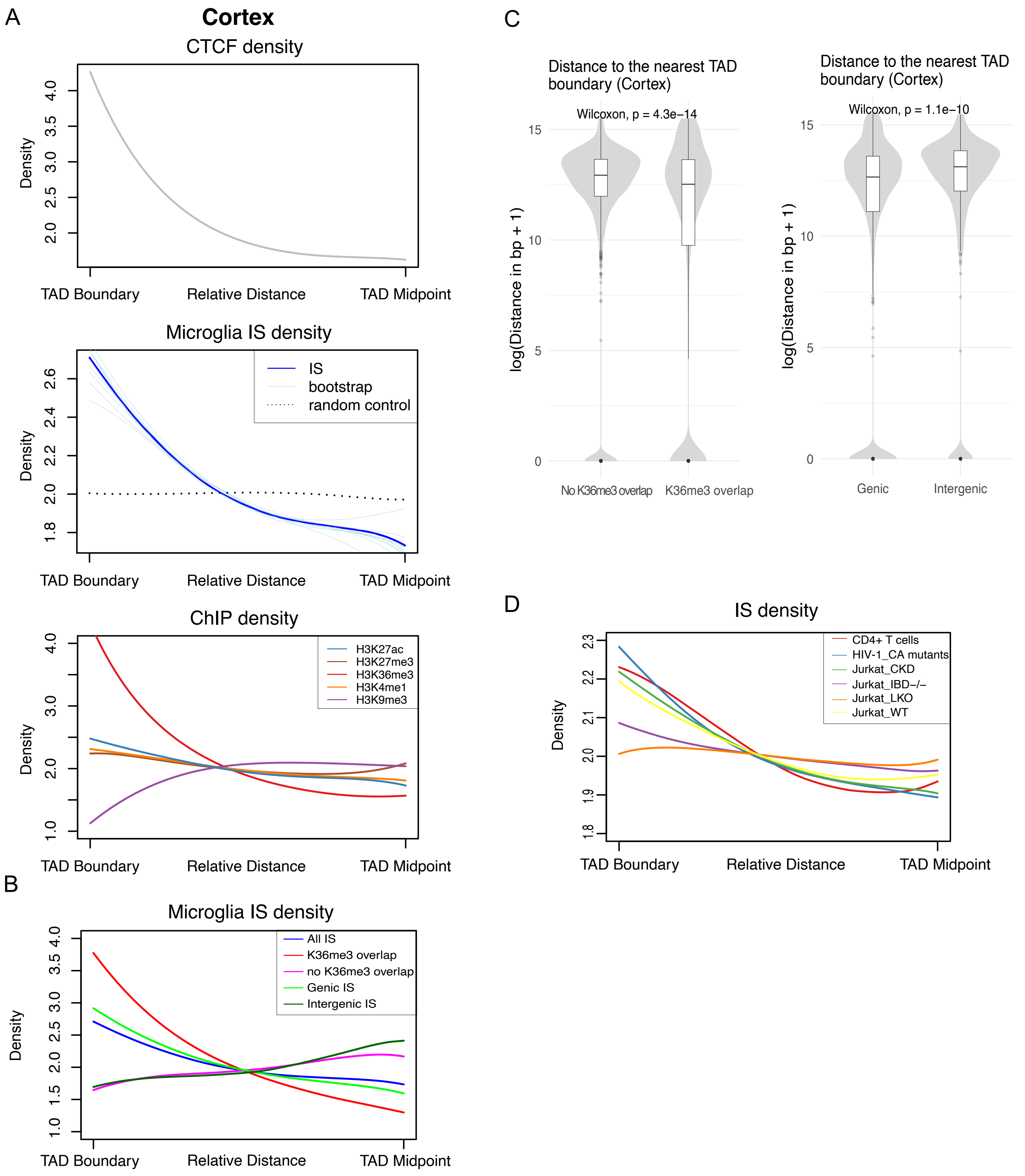


Supplementary Figure S6
